# Attenuated *Salmonella* typhimurium cancer therapy has direct effects on the tumor epithelium in colorectal cancer

**DOI:** 10.1101/741686

**Authors:** Kendle M. Maslowski, Masumi Takahashi, Yumiko Nakanishi, Gillian M. Mackie, Isabel Everard, Alastair Copland, Hirotsugu Oda, Takashi Kanaya, Hiroshi Ohno

## Abstract

Bacterial cancer therapy (BCT) shows great promise for treatment of solid tumors, yet basic mechanisms of bacterial-induced tumor suppression remain undefined. The intestinal epithelium is the natural route of infection for *Salmonella* and thus harbors innate immune defenses which protect against infection. Attenuated strains of *Salmonella enterica* serovar Typhimurium (*S*Tm) have commonly been used in mouse models of BCT, largely with the use of xenograft and orthotopic transplant cancer models. We aimed to better understand the tumor epithelium-targeted mechanisms of BCT by using mouse models of intestinal tumorigenesis and tumor organoid cultures to assess the effectiveness and mechanisms of treatment with aromatase A-deficient *S*Tm (*STm*^Δ*aroA*^). *STm*^Δ*aroA*^ delivered by oral gavage could significantly reduce tumor burden and tumor load in both a colitis-associated colon cancer model (CAC) and in a spontaneous intestinal cancer model, *Apc^min/+^* mice. *STm*^Δ*aroA*^ colonization of tumors caused alterations in transcription of mRNAs associated with epithelial–mesenchymal transition as well as metabolic and cell cycle-related transcripts. Metabolomic analysis of tumors demonstrated alteration in the metabolic environment of *STm*^Δ*aroA*^-treated tumors, suggesting *STm*^Δ*aroA*^ imposes metabolic competition on the tumor. Use of tumor organoid cultures *in vitro* demonstrated that *STm*^Δ*aroA*^ can directly affect the tumor epithelium with alterations in transcripts and metabolites similar to *in vivo*-treated tumors. Thereby, we demonstrate that bacterial cancer therapy is efficacious in autochthonous intestinal cancer models, that BCT imposes metabolic competition, and that BCT has direct effects on the tumor epithelium, which have not previously been appreciated.

**One Sentence Summary:** Attenuated *Salmonella enterica* serovar Typhimurium can home to gastrointestinal tumors and directly affect the tumor epithelium, inducing transcriptional and metabolic changes that lead to reduced tumor burden in mice.

## Introduction

The concept of using bacteria as cancer therapeutics (bacterial cancer therapy, BCT) dates back to the late 1800’s, when the field of BCT was initiated by William Coley (*1*). Even earlier observations of spontaneous tumor regression following infection of patients’ tumors had led Coley to treat cancer patients with injections of bacterial preparations. Despite this early work on BCT there is only one currently in the clinic; Bacillus Calmette–Guérin vaccine (BCG) therapy for superficial bladder cancer (*2*). The advancement of molecular genetics and genetic engineering has enabled bacteria to be effectively attenuated to remove adverse side-effects and engineered to deliver different payloads. With this, there has been a resurgence in interest in BCT in the past 20 years, with many studies showing efficacy of attenuated bacterial treatments in xenograft and orthotopic transplant tumor models, with *Salmonella enterica* serovar Typhimurium (*S*Tm) being by far the most studied (*3–9*). Yet despite this interest, very little is understood about the underlying mechanisms of BCT-mediated tumor suppression, which is holding back its practical application. A few phase 1 trials have been conducted using attenuated *S*Tm strains and showed mixed results in terms of colonization and effect (*10–12*). The *S*Tm strain VNP20009 was used in these trials, and when given i.v. showed poor tumor colonization and induced toxicity (*10*). Another of the trials used intratumoral delivery of VNP20009, and higher tumor colonization was observed without toxicity (*11*). Lack of chemotactic ability of the VNP20009 strain, due to mutation of the *cheY* gene, has been suggested to be a limiting factor to its success. Mouse models have shown *cheY* to be redundant (*13*), while another has shown it to be important, for tumor localization (*14*).

Despite poor translation into the clinic as yet, data from mouse models, as well as Coley’s work and the use of BCG therapy, does suggest promise for BCTs. Attenuated *S*Tm are extremely tumor-tropic—a therapy that can give such tumor tissue selectivity is very desirable and enables further engineering to deliver drugs, immune adjuvants or other anti-tumor agents (*3*). Therefore, further investigation and understanding is warranted. Tumor tissue tropism of attenuated bacteria is thought to be driven by the lack of immune detection within tumors and also the metabolic environment. A benefit of using *S*Tm is that it is a facultative anaerobe, which means it can reside in both anoxic and aerobic regions of tumor. Recruitment and/or retainment of *S*Tm in the tumor environment has been suggested to be driven by availability of metabolites. High levels of aspartate, serine, ribose/galactose and ethanolamine have been shown to play a role in different *in vitro* and *in vivo* cancer models (*14, 15*). The role of chemotaxis is contested, with results suggesting it to be both important and redundant (*13, 14, 16*). Crull *et al* (*16*) hypothesized that tumor invasion *in vivo* is more passive than *in vitro*, as the resulting chemokine and cytokine release upon i.v. or i.p. delivery of *S*Tm would open tumor vasculature enabling delivery of bacteria to the tumor. Therefore, the delivery route and tumor type might account for varying results. It appears that invasion of tumor cells is not necessary for anti-tumor effects, as mutants of invasion genes (eg. SPI-1 and SPI-2) are equally good at invading tumor tissue and inducing tumor regression (*14, 16*). One study showed the formation of *S*Tm biofilms within tumors, which was important for their anti-tumor effects (*17*). It may well be that the tumor model and mode of delivery play a large role in the requirements of *S*Tm, or other bacteria, to home to tissue.

BCT is assumed to exert its effect by eliciting an immune response. Yet there isn’t a clear understanding of immune-mediated events, whether an immune response is elicited to the bacteria, or to tumor cells. It is thought that attenuated *S*Tm, or other bacteria, can survive within tumors because there is poor immune response (*3*). Effective treatment with *S*Tm has been observed in adaptive immune-deficient (nude) mouse models (*8, 18*), suggesting that T cell responses may not be essential for anti-tumor effects. In some studies that have assessed immune infiltrates, only *S*Tm or *E. coli* that were engineered to deliver an immune-stimulatory product elicited a significant immune response (*19, 20*). Other studies have shown induction of an immune response, such as induction of IL-1*β* and TNF-*α* when delivered i.v. (*21, 22*) or increased neutrophil and CD8 T cell recruitment when delivered orally (*9*) to mice bearing subcutaneous tumors (CT26 or MC38 colorectal tumors). So, whilst there is some evidence of changes in tumor immune responses following BCT, it is not yet known whether this is key to the anti-tumor effects of BCT.

The hypothesis that we aimed to explore here is that BCT can have a direct effect on the tumor epithelial cells. Interestingly, the only current BCT in the clinic, BCG used for superficial bladder cancer (*2*), is delivered intravesically directly to the bladder epithelium, which might be important for its success. Studies on BCT so far have utilized xenograft or orthotopic transplant models of cancer, which may not fully model complex disease in patients and therefore not inform on the mechanisms of BCT. Driven by an interest in intestinal epithelial innate immune pathways that protect epithelial cells from *S*Tm infection and tumorigenesis (*24, 25*), we aimed to determine if *S*Tm treatment could be effective for treating autochthonous tumors of the intestine. We reasoned that since *S*Tm naturally infects via the intestine, that intestinal cancer would be a good model to use, and that treating colorectal cancer (CRC) patients by oral delivery of attenuated *S*Tm would be feasible, since oral vaccines for *S*. typhi are widely used and tolerated (*26*). Oral delivery of *S*Tm may also avoid problems of tumor homing and toxicity that have been observed when delivering *S*Tm i.v. to patients (*10*). Using a model of colitis-associated cancer (CAC) and a spontaneous model of intestinal cancer, *Apc^min/+^* mice, we showed that oral delivery of an attenuated *S*Tm reduces tumor burden. Transcriptomic and metabolomic analysis, coupled with use of tumor organoids *in vitro*, demonstrated effects of *S*Tm on the tumor epithelium, and found that metabolic competition was a likely driver of anti-tumor effects.

## Results

### Orally-administered aromatase-A-deficient *S*Tm reduces gastrointestinal tumor burden

We utilized *S*Tm deficient for aromatase A (*STm*^Δ*aroA*^) (UF020 (*27*)) to assess whether BCT could be effective for treating murine autochthonous gastrointestinal tumors (colon or small intestine (SI)). We first induced colon tumors in C57B6/J mice using a well-described model of colitis-associated cancer (CAC), which has 100% penetrance (*25, 28*) (Fig. 1A). After tumor induction, mice were then split into treatment groups, ensuring equivalent colitis severity between groups. Fig. S1 shows weight loss during the AOM/DSS protocol. After recovery from the final dose of DSS, mice were given 5×10^9^ CFU *STm*^Δ*aroA*^, or vehicle control, by oral gavage once per week for 6 weeks. After 6 weeks treatment, tumor burden was assessed and compared to that of control-treated mice and to mice sacrificed at the start of the *STm*^Δ*aroA*^ treatment protocol (denoted D0) (Fig. 1A). Tumor burden and tumor load was significantly decreased in *STm*^ΔaroA^-treated mice, compared to D0 and 6-week control-treated mice (Fig. 1B). This indicates that *STm*^Δ*aroA*^ treatment by oral delivery could reduce existing tumor burden and prevent further tumor development or growth.

**Figure 1:**
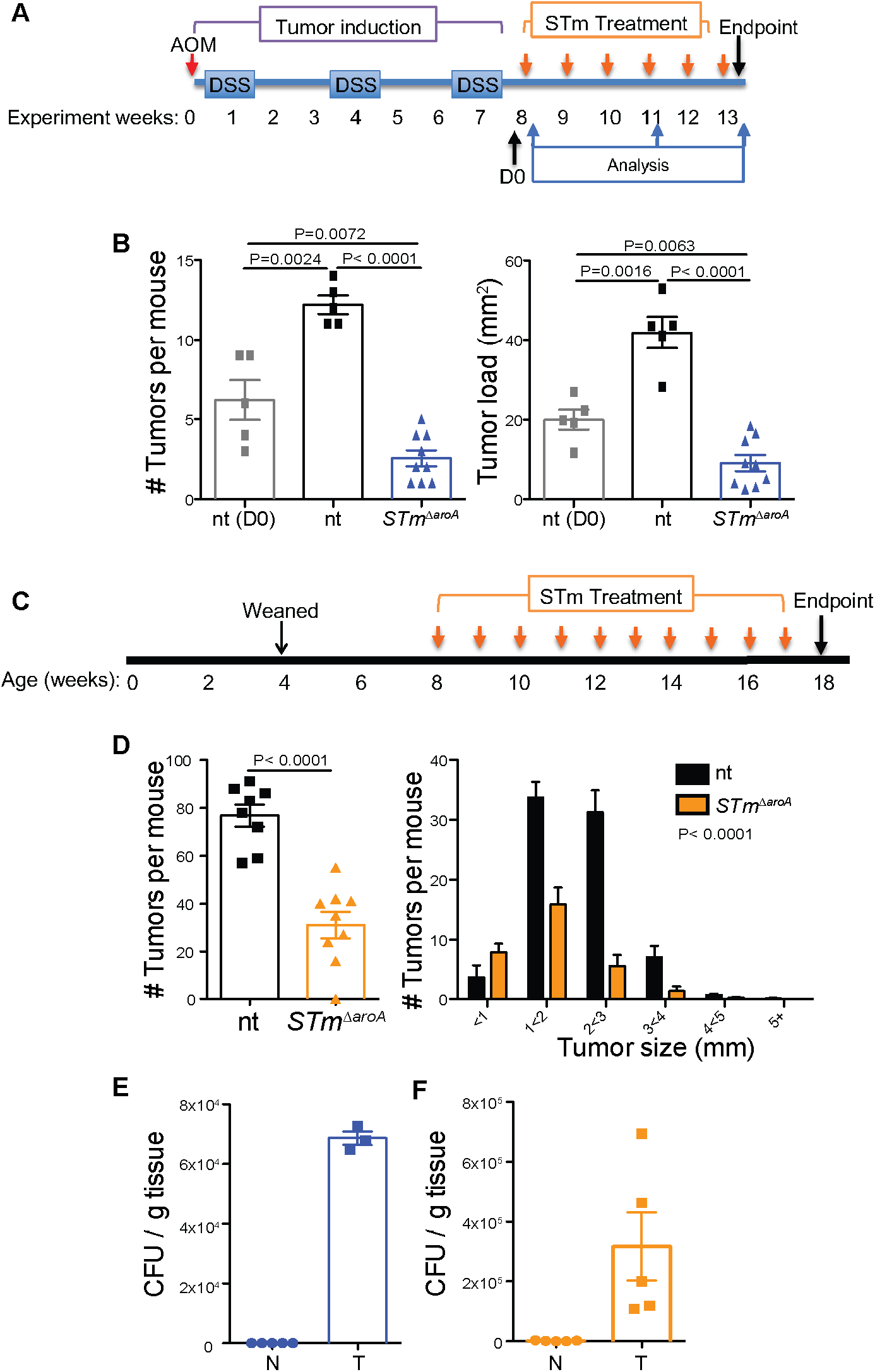
Oral delivery of attenuated *S*Tm reduces intestinal tumor burden. (**A**) Schematic of AOM/DSS induced CAC model and *STm*^Δ*aroA*^ treatment. (**B**) Tumor burden (# tumors / mouse) and tumor load (cumulative tumor size per mouse, mm^2^) in non-treated (nt) and *STm*^Δ*aroA*^-treated mice. N=5 for D0 and nt groups and N=9 for *STm*^Δ*aroA*^-treated mice. Representative of 4 independent experiments. Female mice were used in this experiment, male mice also showed same response. (**C**) Schematic of *Apc^min/+^* mouse *STm*^Δ*aroA*^ treatment. (**D**) Tumor burden and tumor size per mouse in non-treated (nt) and *STm*^Δ*aroA*^-treated mice. Data pooled from 2 independent experiments using both male and female mice, nt N=8 (4F, 4M), *STm*^Δ*aroA*^-treated N=9 (5F, 4M). (**E-F**) colony forming units (CFU) of *STm*^Δ*aroA*^ in normal (N) and tumor (T) tissue from *STm*^Δ*aroA*^ treated mice in the CAC and *Apc^min/+^* models, respectively. One-way ANOVA (B, D) or two-way T-test (D) were used.

Next, we tested *STm*^Δ*aroA*^ treatment in *Apc^min/+^* mice. These mice carry a mutation in the adenomatous polyposis coli gene (*Apc*) which results in multiple intestinal neoplasia (min), serving as a model of human familial adenomatous polyposis (FAP). Though, in mice the *Apc* mutation results largely in small intestinal (SI) neoplasia, and not colonic neoplasia. We treated *Apc^min/+^* mice with 5×10^9^ CFU *STm*^Δ*aroA*^ by oral gavage once per week for 10 weeks, from 8 weeks of age (Fig. 1C). At this age, the SI has already developed a large number of polyps and they continue to grow in size, with mice at 18 weeks showing large well-developed polyps throughout the SI tract. Treatment of *Apc^min/+^* mice with *STm*^Δ*aroA*^ substantially reduced the tumor burden and tumor size (Fig. 1D).

### *STm*^Δ*aroA*^ specifically localize in tumor tissue

Attenuated strains of *S*Tm preferentially home to tumor tissues and not to normal healthy tissues (*4–9*) and hence do not cause overt disease in recipient mice. We confirmed that *STm*^Δ*aroA*^ could be re-isolated from tumor tissue, but not from normal healthy tissue surrounding the tumors in both the CAC and *Apc^min/+^* models (Fig. 1E **and** F, respectively). Further, we checked whether *STm*^Δ*aroA*^ could disseminate to peripheral organs 24 hours after infection of *Apc^min/+^* mice and could find very few or no CFU in the blood, MLN, liver or spleen (Fig. S2).

We next employed scanning electron microscopy (SEM) to view bacterial colonization in greater detail. We found exceptionally large colonies of *STm*^Δ*aroA*^ within the tumor mass just 24 hrs after administration (Fig. 2, see insets). The large size of the bundles suggest that they were rapidly dividing within the tumor extracellular spaces. We could also find instances of single or a few bacteria (Fig. 2, red arrows). No bacteria could be observed in non-treated mice (Fig. S3), suggesting that normal microbiota are not able to penetrate tumor tissue to form mass colonies as observed with the *STm*^Δ*aroA*^. Immunohistochemistry detecting mCherry-expressing *STm*^Δ*aroA*^ further supports the SEM data showing large colonies of *STm*^Δ*aroA*^ (Fig. S3).

**Figure 2:**
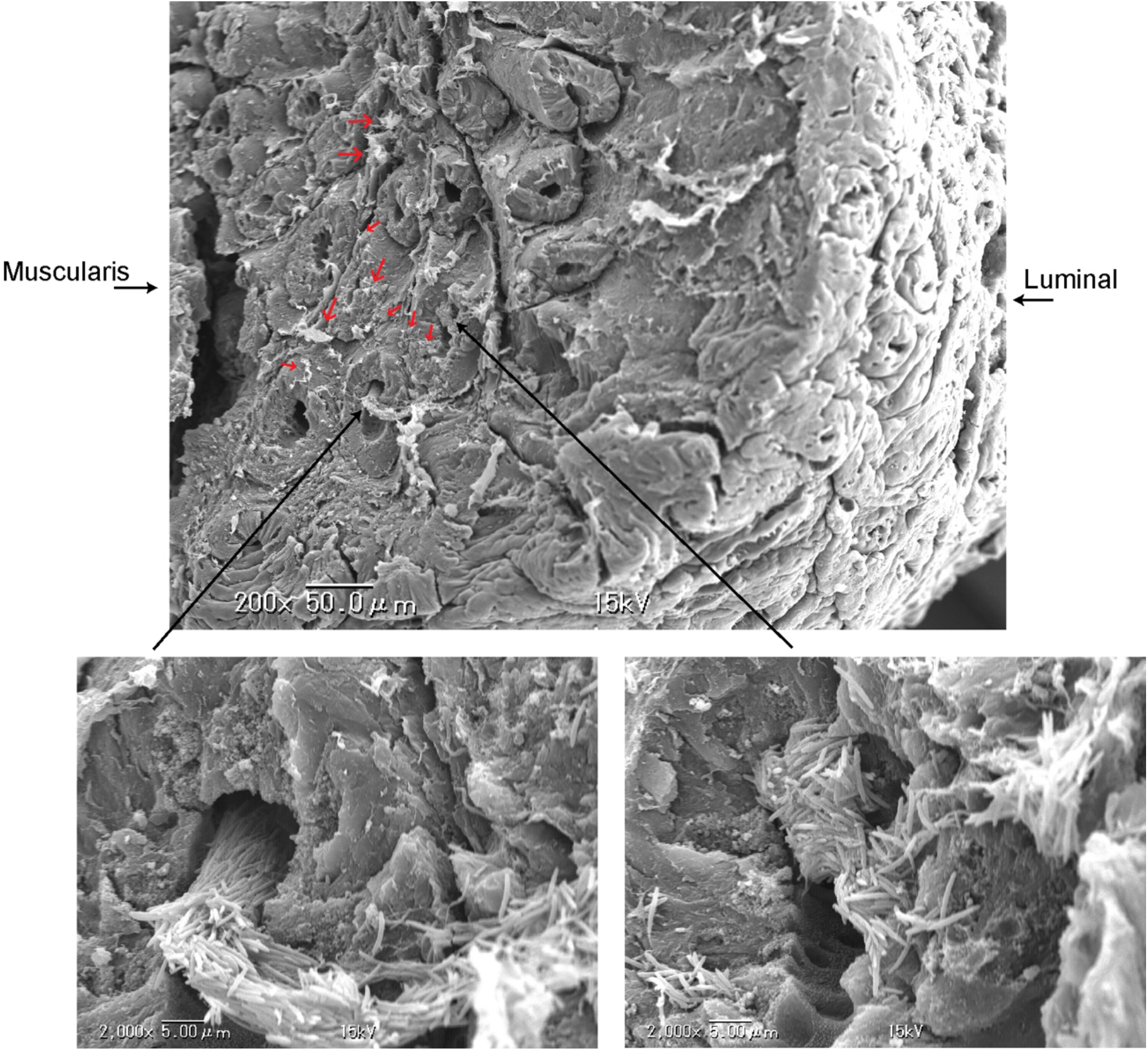
Scanning electron microscopy of *STm*^Δ*aroA*^-treated tumors. Mice bearing CAC colon tumors were given *STm*^Δ*aroA*^ or control vehicle by oral gavage and tissues were taken 24 hours later. Whole sections of colon with tumors were prepared for SEM by glutaraldehyde fixation, dehydration and freeze drying. Tumors were cut on the sagittal plane and mounted for platinum-coating and SEM imaging. Top image shows lower magnification view of a tumor area. Luminal side indicates the top of the tumor that was facing the intestinal lumen and muscularis side indicates the inner side of tumor reaching the lamina propria and muscularis mucosa. Small red arrows indicate small *STm*^Δ*aroA*^ colonies or individual bacteria. Large black arrows indicate areas shown in higher magnification shown below.

### *STm*^Δ*aroA*^ alters the transcriptional landscape of tumors

Next, to gain an understanding of the differences between non-treated and *STm*^Δ*aroA*^-treated tumors, we performed RNA sequencing on RNA isolated from whole tumor homogenates. Tumor (T) and adjacent normal (N) tissue was dissected from AOM/DSS-induced CAC-bearing mice after 4 weeks treatment. First, we identified the transcripts that were differentially regulated between N and T tissue in the non-treated and *STm*^Δ*aroA*^-treated groups. Fig 3A shows the number of overlapping and unique genes for each treatment. These differentially-expressed genes (DEGs) were then analyzed by gene ontology (GO) analysis using DAVID (*29, 30*), revealing terms enriched in either the non-treated tumors or in the treated tumors, which intriguingly were vastly different (Fig. 3B). As expected, non-treated tumors exhibited enrichment of mRNAs involved in cell cycle processes, mitosis, cell division, DNA repair etc., whereas, *STm*^Δ*aroA*^-treated tumors displayed enrichment of mRNAs for processes involving regulation of mesenchymal cell proliferation, mesenchymal-epithelial cell signaling, as well as regulation of blood vessel development, amongst other pathways. (Fig. 3B, Fig. S4).

**Figure 3:**
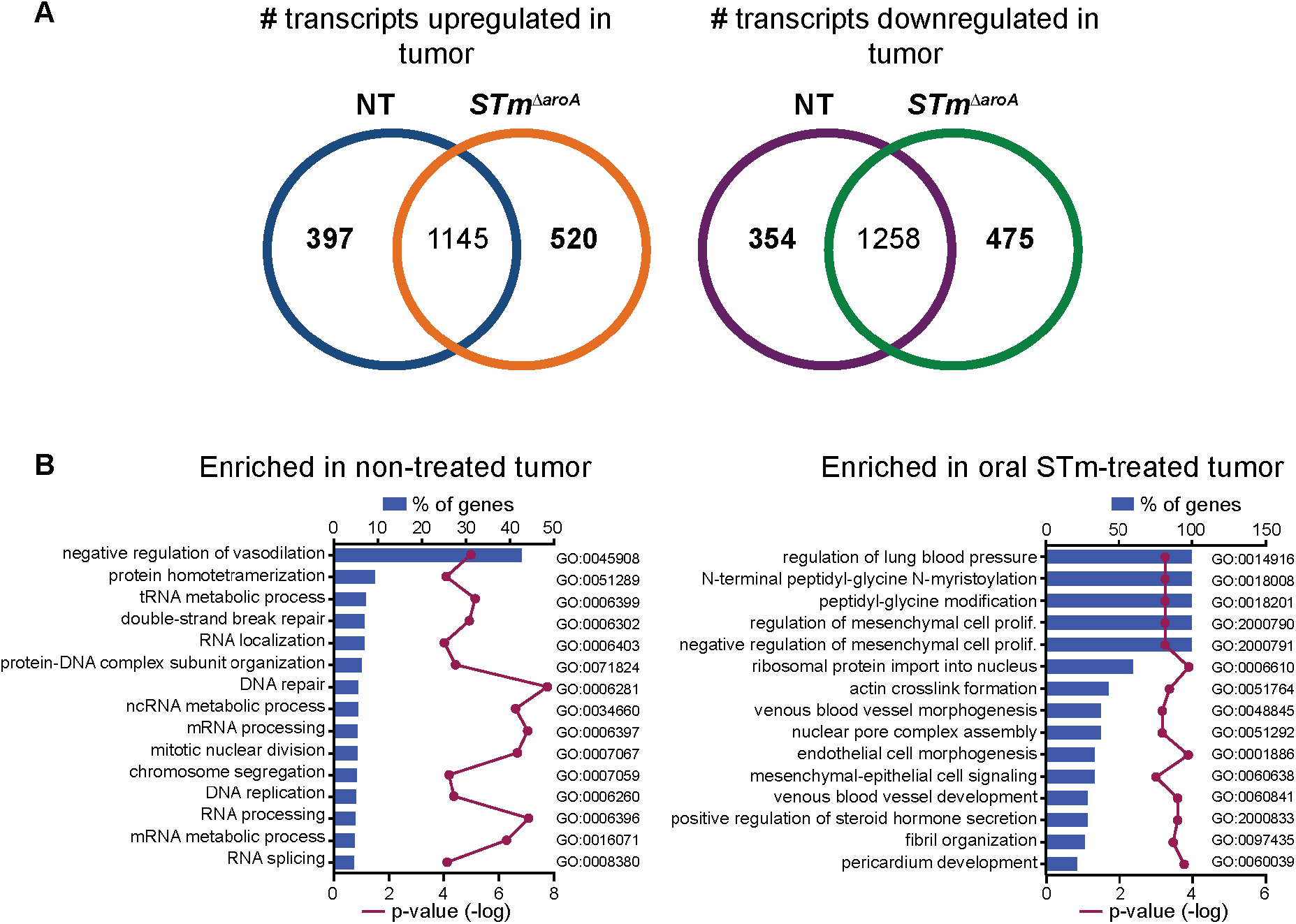
*STm*^Δ*aroA*^ treatment alters the transcriptional landscape of tumors. Normal and tumor tissue were dissected from CAC-bearing mice after 4 weeks of *STm*^Δ*aroA*^ or control treatment and RNA isolated and used for RNA sequencing analysis. (**A**) # of transcripts upregulated or downregulated in tumor compared to normal tissue, with overlapping and unique transcripts depicted. (**B**) Differentially expressed genes (DEGs) in tumors compared to normal tissue for each treatment identified in (A) were compared by GO analysis. Data represents the % of genes of a given pathway that are enriched in either non-treated or treated tumors, with -log p-value.

Several genes involved in DNA repair, DNA damage response, RNA synthesis and epithelial-mesenchymal transition were significantly reduced following *STm*^Δ*aroA*^ treatment (Fig. S4), which suggest a change in rate of cell proliferation. We decided to concentrate on stem cell, EMT and metabolism-related genes, and confirmed a selection of targets by qPCR in independent experiments. As previously reported, transcripts for epithelial stem cells, proliferation or epithelial-to-mesenchymal transition-related processes, including *Lgr5* (leucine-rich repeat-containing G-protein coupled receptor), *Smoc2* (SPARC related modular calcium binding 2), *Vim* (Vimentin), *Ccnd1* (Cyclin D1) and *Pdk4* (pyruvate dehydrogenase kinase 4) (*31–37*) were increased in tumor tissue when compared to normal tissue (Fig. 4A). Strikingly, these transcripts were largely decreased following *STm*^Δ*aroA*^ treatment (Fig. 4A). We confirmed these mRNA changes in the *Apc^min/+^* model, comparing tumor tissue from non-treated and *STm*^Δ*aroA*^ treatment. In line with results from the CAC model, *STm*^Δ*aroA*^ treatment altered the transcriptional levels of the above-mentioned genes and additional EMT-related genes *Twist* and *Snail* (Fig. 4B). Loss of E-cadherin protein at the cell-cell junctions is an important feature of EMT, therefore we checked E-cadherin protein expression and localization by immunofluorescent staining of sections taken from CAC tumor-bearing mice. Non-treated tumor sections showed very little E-cadherin protein; and where it was present it was cytosolic staining (Fig. 4C). In contrast, tumors from *STm*^Δ*aroA*^ treated mice showed significant levels of cell membrane localized E-cadherin (Fig. 4C). Thus it appears that *STm*^Δ*aroA*^ treatment diminishes tumors and restores epithelial identity. This likely reflects that *STm*^Δ*aroA*^ treatment is having a direct effect on tumor cells as well as the fact that tumors are diminished in size and normal epithelium is restored.

**Figure 4:**
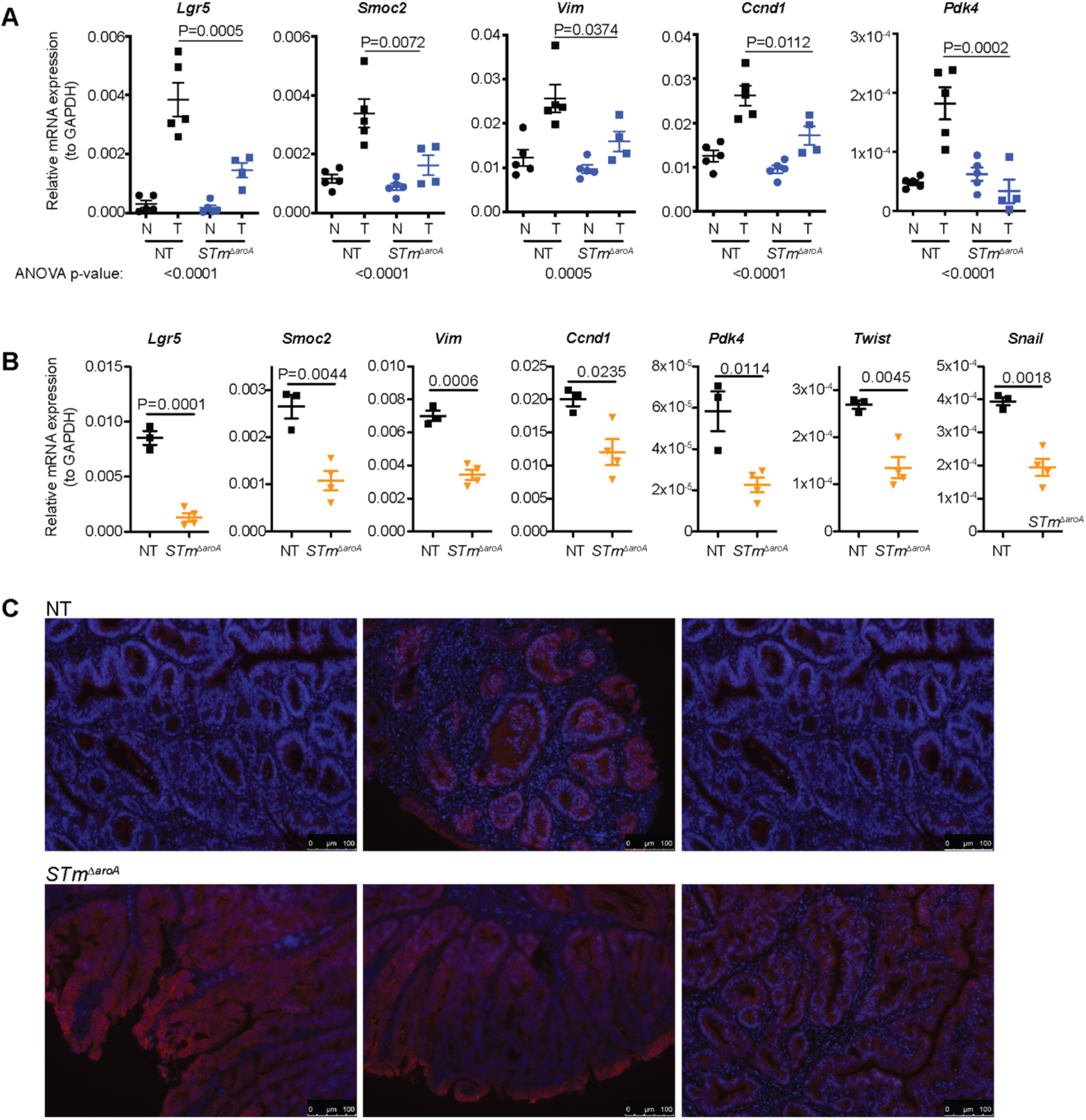
Altered tumor phenotype in *STm*^Δ*aroA*^ treated mice. (**A**) qPCR confirmation of genes identified (or pathway related) by RNAseq in CAC-tumor bearing mice after 6 weeks treatment. Non-treated (NT), Salmonella treated (*STm*^Δ*aroA*^), normal tissue = N, tumor tissue = T. Data representative of 4 independent experiments. One-way ANOVA with Turkey’s multiple comparisons test was conducted. ANOVA P-values are indicated below the graphs and individual post-test comparing T from each treatment is shown on the graphs. (**B**) Analysis of indicated transcripts in *Apc^min/+^* tumor tissue after 10 weeks treatment. Data representative of 2 independent experiments. NT N = 3, *STm*^Δ*aroA*^ N=4. Unpaired two-way T-tests were used. (**C**) Immunofluorescence of E-cadherin (red) counterstained with DAPI (blue) in NT and *STm*^Δ*aroA*^-treated CAC mice.

### *STm*^Δ*aroA*^ alters the metabolic environment of tumors

Previous studies have demonstrated that that BCT can affect tumor growth by utilizing excess nutrients, such as ethanolamine (*14*), or are attracted to tumors due to high levels of metabolites, such as ribose or leucine (*15*). Our observation of large intra-tumoral, extracellular *STm*^Δ*aroA*^ colonies led us to question whether the tumor metabolome would be altered following treatment, as the bacteria would be competing for metabolites within the tumor environment. Tumor and normal tissue from non-treated or *STm*^Δ*aroA*^-treated CAC-tumor bearing mice after 6 weeks or 24 hours treatment were analyzed by GC-MS for polar-metabolites. Unit variance (UV) scaled GC-MS data was analyzed, and Orthogonal Partial Least Squares-Discriminant Alanysis (OPLS-DA) plots revealed a separation between non-treated and treated tumors after 6 weeks and also within 24 hours (Fig. 5A, B respectively) (6 weeks treatment in vivo: R^2^ = 0.99; Q^2^ = 0.52; 24 hours treatment in vivo: R^2^ = 0.99; Q^2^ = 0.67). We performed pathway analysis on metabolites with a variable importance on the projection (VIP) score greater than 1 using MetaboAnalyst 3.0 (*3, 4*) (Tables S1 and S2 show the complete list). Common pathways affected by *STm*^Δ*aroA*^ treatment at both timepoints (6 hours and 24 hrs) included glycine, serine and threonine metabolism, arginine and proline metabolism and citric acid cycle, amongst others (Fig. 5C,D). As previously described (*40*), many metabolites, and particularly amino acids and tricarboxylic acid (TCA) cycle intermediates, are increased in tumor tissue compared to normal tissue (Fig. 5E, Fig. S5 & 6). This likely reflects the increased energy and anabolic requirements of tumors. Strikingly, many metabolites were decreased following *STm*^Δ*aroA*^ treatment. Fig 5E shows metabolites detected in glycolysis and the TCA cycle, as well as amino acids. Perhaps not surprisingly, glucose was significantly reduced in *STm*^Δ*aroA*^ treated tumors (Fig 5E). Other glycolysis intermediates were only mildly affected, while several TCA cycle intermediates (citrate, succinate, fumarate and malate) were reduced following treatment. (Fig 5E). Furthermore, several amino acids, which can feed different parts of the TCA cycle were reduced, including glutamate (glutamine was not detected) (Fig 5E), which is an important fuel for glutaminolysis for which tumors can be quite dependent (*41*).

**Figure 5:**
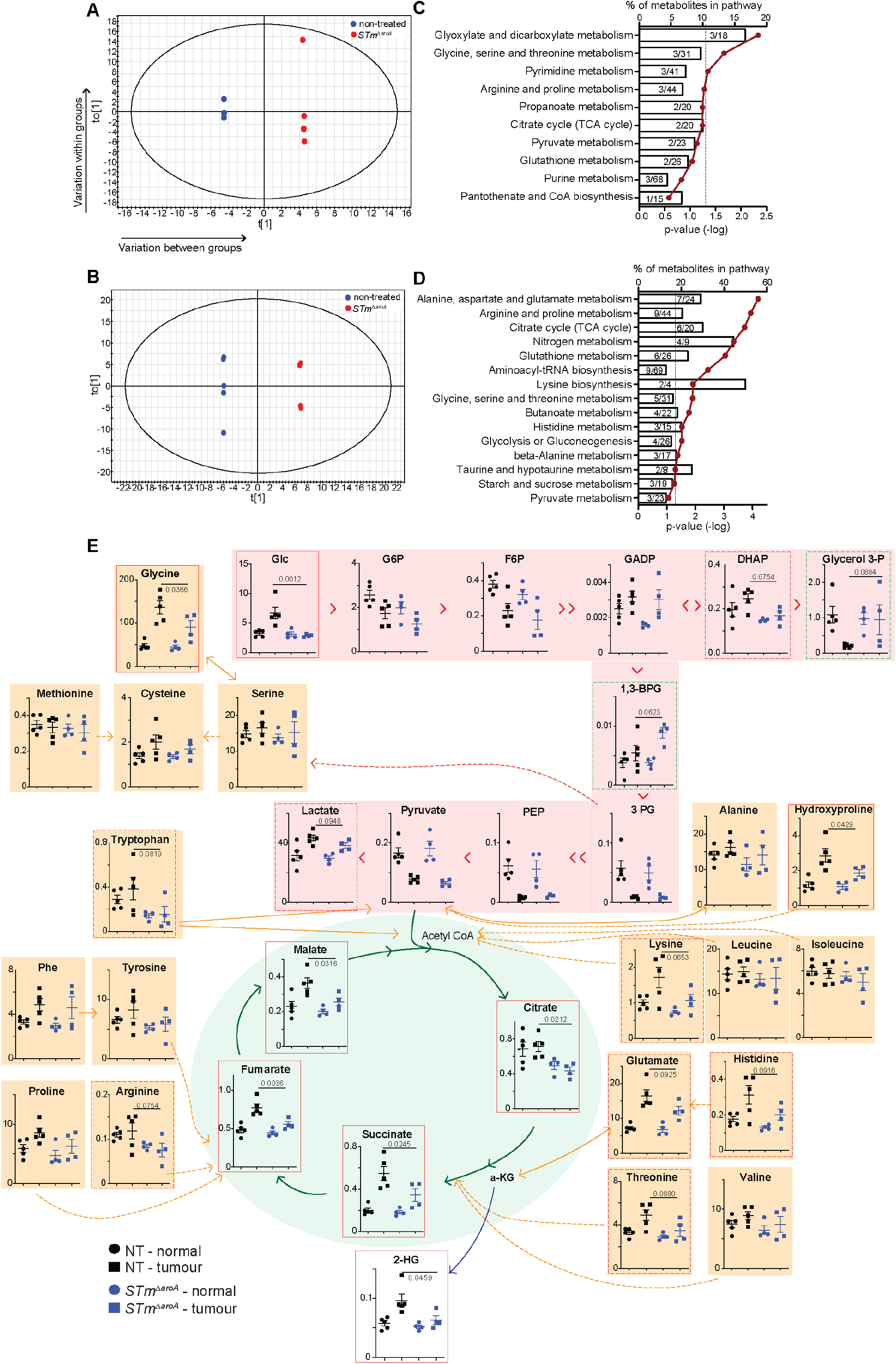
*STm*^Δ*aroA*^ treatment alters the metabolic environment of tumors. Tumor metabolites were assessed by GC-MS. OPLS analysis of metabolites comparing non-treated (NT) and *STm*^Δ*aroA*^ treated tumors after (**A**) 6 weeks and (**B**) 24 hours treatment. All metabolites significantly different between *STm*^Δ*aroA*^ treated and non-treated tumors (VIP score>1) were submitted to pathway analysis (MetaboAnalyst). Pathway analysis for 6 weeks *STm*^Δ*aroA*^ treatment (**C**) and 24 hours treatment (**D**), represented as the percent of metabolites in a pathway that were altered, against p-value (-log). (**E**) Metabolites detected from glycolysis (pink shading) and TCA cycle (green shading), and amino acids (orange shading), with interrelationships depicted. X-axis are nmol/g. One-way ANOVA was performed with Bonferroni multiple comparisons test, P-values shown are the multiple comparison statistic.

Other important oncometabolites were also affected. The polyamine synthesis pathway appears affected at both 24 hours and 6 weeks post-treatment. Ornithine was reduced in *STm*^Δ*aroA*^ treated tumors 24 hours and 6 weeks post-treatment, while putrescine was significantly affected after 6 weeks treatment and spermidine after 24 hours (Fig. S5 & 6). 2-hydroxyglutarate (2-HG) can accumulate in tumors due to mutant isocitrate dehydrogenase 1/2 (IDH2) activity, converting *α*-ketoglutarate (*α*-KG) to 2-HG, which can inhibit *α*-KG-dependent dioxygenases, leading to increased histone and DNA methylation (*41*). We saw increased 2-HG in CAC tumors, and this was decreased after 24 hours treatment (Fig. 5E). Thus, several important fuel sources, metabolic intermediates and oncometabolites are decreased following *STm*^Δ*aroA*^ treatment, assumedly through metabolic competition of the bacteria on the tumor cells.

### *STm*^Δ*aroA*^ directly affects tumor epithelium

Our initial hypothesis was that BCT would have a direct effect on tumor epithelium. The effects that we have described so far on tumor stem cell markers, an increase in epithelial identity, and a change in the tumor metabolome in *STm*^Δ*aroA*^ treated tumors, suggests an impact of *STm*^Δ*aroA*^ treatment on tumor cells. To directly test our hypothesis that *STm*^Δ*aroA*^ treatment has a direct effect on tumor epithelium, we utilized tumor 3D organoid cultures. We generated organoid lines from CAC-induced colorectal tumors and from *Apc^min/+^* SI and colonic tumors. Representative images of organoid appearance is shown in Fig. S8A-C. Tumor organoids were infected with *STm*^Δ*aroA*^ by inoculating the culture medium (1×10^8^ CFU). *STm*^Δ*aroA*^ were able to invade the Matrigel and infect the organoids (Fig. 6A&B). After 2 hrs infection, the culture medium was washed off and fresh medium containing gentamycin was added, so only bacteria that had infected organoids could grow, preventing any effects purely from bacterial over-growth. Organoids were then collected for analysis 24 hours after the initial infection. CFU analysis was performed to determine bacterial burden. We observe in the order of 1×10^5^ CFU (Fig. 6B) per well, which contains around 1×10^6^ cells within the organoid structures. Importantly, treatment of organoids with *STm*^Δ*aroA*^ could recapitulate effects on gene expression seen *in vivo*, with a substantial reduction in transcripts for *Lgr5*, *Smoc2* and *Vim* in both CAC-derived and *Apc^min/+^*-derived tumor organoids, and Pdk4 in *Apc^min/+^* organoids (expression was very low in CAC organoids) (Fig. 6C and D). These results provide evidence that *STm*^Δ*aroA*^ treatment can directly affect the tumor cells, independent of effects on other systems/cell types such as the immune system. Furthermore, analysis of the organoid metabolome demonstrated separation of non-treated and treated organoids by OPLS analysis (Fig. 6E). Taking all metabolites with a VIP score >1 (Table S3) and analyzing by MetaboAnalyst revealed similarly affected metabolic pathways following *in vitro STm^ΔaroA^* treatment as for *in vivo* treatment, with amino acid metabolism pathways, TCA cycle and glycolysis being altered (Fig. 6F, Fig. S7). These data suggest that bacterial colonization imposes direct metabolic competition.

**Figure 6:**
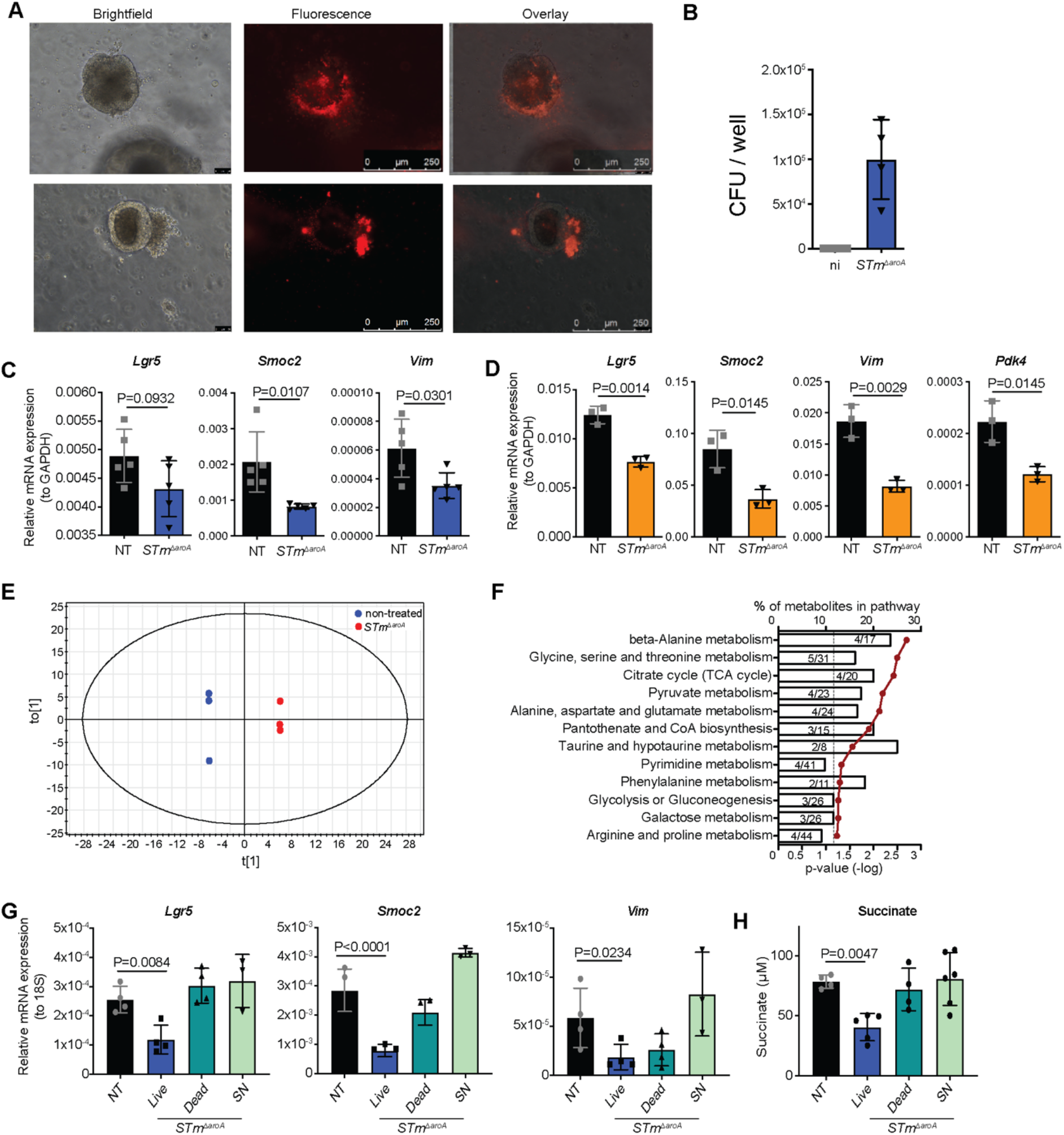
*In vitro* treatment of tumor-derived organoids with *STm*^Δ*aroA*^. Tumor organoids derived from CAC-induced tumors and *Apc^min/+^* tumors were established and infected with mCherry-expressing *STm*^Δ*aroA*^ for 24 hrs. (**A**) Brightfield and fluorescent microscope images of organoids within matrigel after 24 hrs infection, shows association of *STm*^Δ*aroA*^ with tumor organoids and the appearance of bursting. (**B**) CFU of *STm*^Δ*aroA*^ per well after 24 hrs infection. qPCR analysis of the indicated transcripts in (**C**) CAC-derived and (**D**) *Apc^min/+^* tumor organoids. Representative of >3 independent experiments with 4 independently-derived organoid lines, with between 3 and 5 technical replicates per experiment. Tumor organoid metabolites were assessed by GC-MS, (**E**) OPLS-DA analysis and (**F**) pathway analysis of metabolites with a VIP score>1. (**G**) CAC-derived tumor organoids were cultured with live or heat-killed (dead) *STm*^Δ*aroA*^, or with supernatant (SN) of *STm*^Δ*aroA*^ grown in organoid culture medium and the indicated mRNA’s analyzed by qPCR. (**H**) analysis of succinate levels in tumor organoids treated as described in (**G**). Individual two-way T-tests (**C**&**D**) or Kruskal-Wallis tests (**G-H**) were performed.

To further dissect whether live bacteria are required to mediate the observed effects of *STm*^Δ*aroA*^ on tumor organoids, or whether the presence of heat-killed bacteria or just the bacterial supernatant would be sufficient, we compared treatment of tumor organoids with live *STm*^Δ*aroA*^, heat-killed *STm*^Δ*aroA*^ or *STm*^Δ*aroA*^ supernatant (SN, prepared using 10 kDa exclusion columns). Live bacteria had the strongest effect on reducing stem cell and EMT marker expression (Fig. 6G). Heat-killed bacteria induced a slight reduction in *Smoc2* and *Vim*, while *STm*^Δ*aroA*^ SN had no effect (Fig. 6G), suggesting that secreted products from bacteria are not exerting these anti-tumor effects, and bacterial ligands (eg LPS) are also not driving these responses (note 10kDa filters exclude LPS, Fig. S8D). Furthermore, succinate, one of the metabolites identified as being reduced by *STm*^Δ*aroA*^ treatment *in vivo*, was assayed and only live *STm*^Δ*aroA*^ could induce a reduction (Fig. 6H), thus further supporting the idea that *STm*^Δ*aroA*^ directly impose metabolic competition.

## Discussion

Here, we present data showing that BCT can be efficacious in autochthonous models of intestinal cancer. Oral delivery of *STm*^Δ*aroA*^ to colonic or SI tumor-bearing mice induced a reduction in tumor number and size. While the concept of using bacteria as cancer therapeutic agents is by no means novel, attempts at mechanistic understanding using *in situ* cancer models has not been attempted before. Previous studies have utilized orthotopic or xenograft transplant tumor models, which may not fully recapitulate complex tumor environments in spontaneously formed tumors. Therefore, to start to unpick mechanisms of BCT action, using tumor models that more faithfully recapitulate tumor development may be necessary. Apart from one BCT in the clinic (BCG used for superficial bladder cancer), trials using attenuated *S*Tm have not been successful. In a phase 1 trials, giving heavily attenuated *S*Tm intravenously resulted in poor tumor localization (*10*), whereas another small trial giving bacteria by intratumoral injection had better tumor localization (*11*). BCG therapy is given directly onto the bladder epithelium. This suggests that the future of BCT may be where it can be applied more locally. Oral delivery of an attenuated *S*Tm would be feasible, and enable delivery directly to colonic tumors, avoiding toxicity overserved with i.v.-delivered bacteria. Proof-of principle on tolerance and safety of such treatment can be seen with *S.* Typhi vaccination (*26*).

Previous studies have shown that *S*Tm genes involved in ethanolamine catabolism were essential for *S*Tm survival within tumors (*14*), and that *S*Tm utilize nutrient sensing pathways to localize to tumors (*15*). We demonstrate that *STm*^Δ*aroA*^ form large intra-tumoral colonies and drastically re-shape the tumor metabolome within 24 hours. Multiple metabolic pathways were affected by *STm*^Δ*aroA*^ treatment, which would impose strong metabolic pressure on tumors cells, which would possibly make it more difficult for tumors to switch from one pathway to another to meet energy and anabolic requirements. Importantly, these biochemical effects were not seen in surrounded normal tissue. Drugging tumor metabolism is an important avenue for cancer therapy, with standard chemotherapies taking advantage of metabolic weaknesses (*41*). However, not all tumors are susceptible, and side-effects from inhibiting all fast-dividing cells limits metabolic inhibitor usage. Further, some tumors are able to metabolically adapt if one pathway is blocked (*41*). BCT may therefore be an avenue for introducing metabolic competition, coupling this with other metabolic inhibitors may enable lower doses of these drugs that may otherwise cause severe side-effects.

The data presented here shows bulk metabolites from tumors that contain *STm*^Δ*aroA*^, so we can’t decipher which metabolites are bacterial-derived and which are host-derived, but the observed decrease in many metabolites from multiple pathways implies that *STm*^Δ*aroA*^ utilize tumor-derived metabolites. Wildtype *S*Tm have been reported to utilize succinate and lactate within the intestinal environment (*42, 43*), and we found a reduction of succinate *in vitro* only when live *STm*^Δ*aroA*^ were present, suggesting active use of tumor metabolites by *STm*^Δ*aroA*^. It is possible that the metabolic competition also induces a shift in the tumor metabolism. We observed changes in *Pdk4*, a metabolic gene involved in inhibiting pyruvate entering into the TCA cycle. Down-regulation of this following *STm*^Δ*aroA*^ may represent the tumor altering metabolism to reflect the change in available metabolites, or could be a result of reduction of other metabolites such as succinate, which would cause less HIF1*α* stabilization and reduced phosphorylation and activation of *Pdk4*.

Importantly, we have shown here that *STm*^Δ*aroA*^ can directly affect tumor cells, a feature that hasn’t previously been appreciated. That *STm*^Δ*aroA*^ reduced stem cell markers is promising as remaining tumor stem cells are able to form relapsing disease. Whether the effect of *STm*^Δ*aroA*^ on stem cells marker expression was due to *STm*^Δ*aroA*^ being more tropic for stem cells or that stem cells are more susceptible to metabolic pressure isn’t yet clear. There are likely numerous concurrent adaptations occurring, and the data presented here provide some new suggestions of how BCT may be effective.

Several groups are taking the approach to BCT to engineer the bacteria to deliver drugs or other compounds that can further promote tumor death or immune clearance (*19, 20, 44–47*). Given that bacteria home specifically to tumors, they are the ideal device to use to ensure tumor-specific drug targeting (*3*). The data we present here shows that BCT does induce tumor regression in autochthonous models of cancer, and we show strong effects on the tumor metabolome and transcriptome. Though it is apparent that *STm*^Δ*aroA*^ alone does not cure the mice, so further engineering of the bacteria, and/or co-therapies, are required. By understanding the mechanisms of action we could further improve the engineering of bacteria for BCT, for example delivering an engineered bacteria that can better utilize metabolites. Or unpicking how BCT affect stem cell transcripts may lead to clues on how to better target tumor stem cells. Furthermore, rational selection of tumor types and stages to be targeted, type of bacteria and attenuations, and delivery route are all likely to be important for the success of BCT. This paper demonstrates that CRC is an excellent candidate for targeting with BCT via oral delivery of attenuated *S*Tm.

## Materials and Methods

### Animals

C57B/6J mice were purchased from JAX or CLEA (Japan). *Apc^min/+^* mice were purchased from JAX (https://www.jax.org/strain/002020). Animal breeding was conducted under specific pathogen-free conditions at the RIKEN Center for Integrative Medical Sciences Animal Facility and infection experiments were conducted at the conventional facility at Yokohama City University. 12 hr day/night cycles (7am-7pm in both facilities) and chow and water were fed *ad libitum*. All animal experiments were approved by the Institutional Animal Care and Use Committees of RIKEN Yokohama Branch and Yokohama City University.

### Colitis-associated cancer model

C57B6/J mice were purchased at 6 weeks of age and allowed settle into the mouse facility for 1-2 weeks. Starting at 7 to 8 weeks of age, mice were given one i.p. injection of AOM (10 mg/kg, Sigma) in the afternoon. The following day, mice were given DSS (1%, MW 35 000-50 000, MP Biochemicals) in the drinking water for 5 days, followed by 16 days normal water. 2 more doses of DSS (1%) were given for 5 day periods. After the 3^rd^ DSS dose, mice recovered for 1 to 2 weeks before starting the *STm*^Δ*aroA*^ treatment.

### *Apc^min/+^* model

Wild-type C57B/6J female mice were bred with heterozygote *Apc^min/+^* males. Litters were weaned at 4 weeks of age and genotype confirmed. *Apc^min/+^* mice (male and female) were transferred to the experimental facility at 6 weeks of age, and *STm*^Δ*aroA*^ treatment started at 8 weeks. At the end of the protocol, genotypes were re-confirmed.

### *STm*^Δ*aroA*^ treatment

UF020 *Salmonella* typhimurium strain lacking aromatase A (*27*) was grown overnight in LB broth with ampicillin (200 ug/mL). In the morning a 1:20 subculture was grown in LB broth (no ampicillin) to an OD_600_ of 0.7 to 0.8 OD_600_ = 5×10^5^ CFU/mL). Cultures were centrifuged (4000 rpm, 8 mins, room temperature) and washed twice in PBS before resuspending at 5×10^9^ CFU per 100 uL. This was transferred to the animal facility within 30 minutes and mice were delivered 100 uL by oral gavage at the indicated time points.

### Tissue isolation and analysis

At the end of the *STm*^Δ*aroA*^ treatment, mice were killed and colons (CAC) or SI (*Apc^min/+^*) were dissected out, cut longitudinally and washed thoroughly in cold PBS. Tumors were counted under a stereo microscope using an eyepiece graticule with 10×10 mm grid. Tumors and normal tissue were then micro-dissected and either snap frozen in liquid N_2_ for later analysis or used for assessing bacterial colonization (CFU).

### Colony forming unit analysis

Tissue was homogenized in PBS containing 0.1% triton-X using a hand-held homogenizer. After serial dilution, 50 uL was plated onto LB agar plates containing ampicillin (200 ug/mL) and incubated at 37°C overnight. Colonies were counted and CFU calculated per gram of tissue.

### Scanning electron microscopy

Colons were dissected out, cut longitudinally and thoroughly washed in PBS. Sections around 1 cm were cut and fixed in 2.5% glutaraldehyde in 0.1% phosphate buffer for 24 hours at 4°C. Tissue was then dehydrated in increasing concentration of ethanol (50% -> 70% -> 80% -> 90% -> 95%-> 100% -> 100%) for 15 minutes each, then tissues substituted with t-butyl alcohol followed by freeze drying. Before mounting samples, tumors were cut on the sagittal plane to reveal the tumor core then mounted onto aluminum stubs and were metal-coated using a magnetron sputter (MSP-1S; Vacuum Device), and examined by scanning electron microscopy (VE-7800; Keyence).

### RNA isolation

RNA was isolated using Qiagen RNeasy Mini kits, as per manufacturer instructions. Briefly, tissue stored in −80°C was placed in buffer RLT + *β*-mercaptoethanol and homogenized by bead beating. Lysates were then processed as recommended, with the optional DNase digest. RNA was quantified using a DeNovix DS-11.

### cDNA library prep and RNA sequencing

RNA integrity was assessed by Agilent Bioanalyzer before proceeding RIN values of 8 or greater were used. 2 ug of RNA was used to prepare a cDNA library using TruSeq RNA Library Prep Kit v2 (Illumina). Sequencing was performed on an Illumina HiSeq 1500 System in a 1×50 bp single read mode. Sequenced reads were mapped against the mouse reference genome (mm10) using TopHat (*48*), and gene expression was quantified by Cufflinks (*49*). Gene ontology enrichment analysis was performed using DAVID (*29, 30*). The original RNAseq data is uploaded and available online (Gene Expression Omnibus: GSE136029).

### cDNA prep and qPCR

cDNA was prepared using standard oligoDT and M-MLV reverse transcriptase. Quantitative real-time PCR was performed with the LightCycler® 480 Real-Time PCR System (Roche, Switzerland) and SYBR Premix Ex Taq (Takara). Gene-specific primers (Eurofins Genomics, Japan) are listed in Table 1.

**Table 1:**
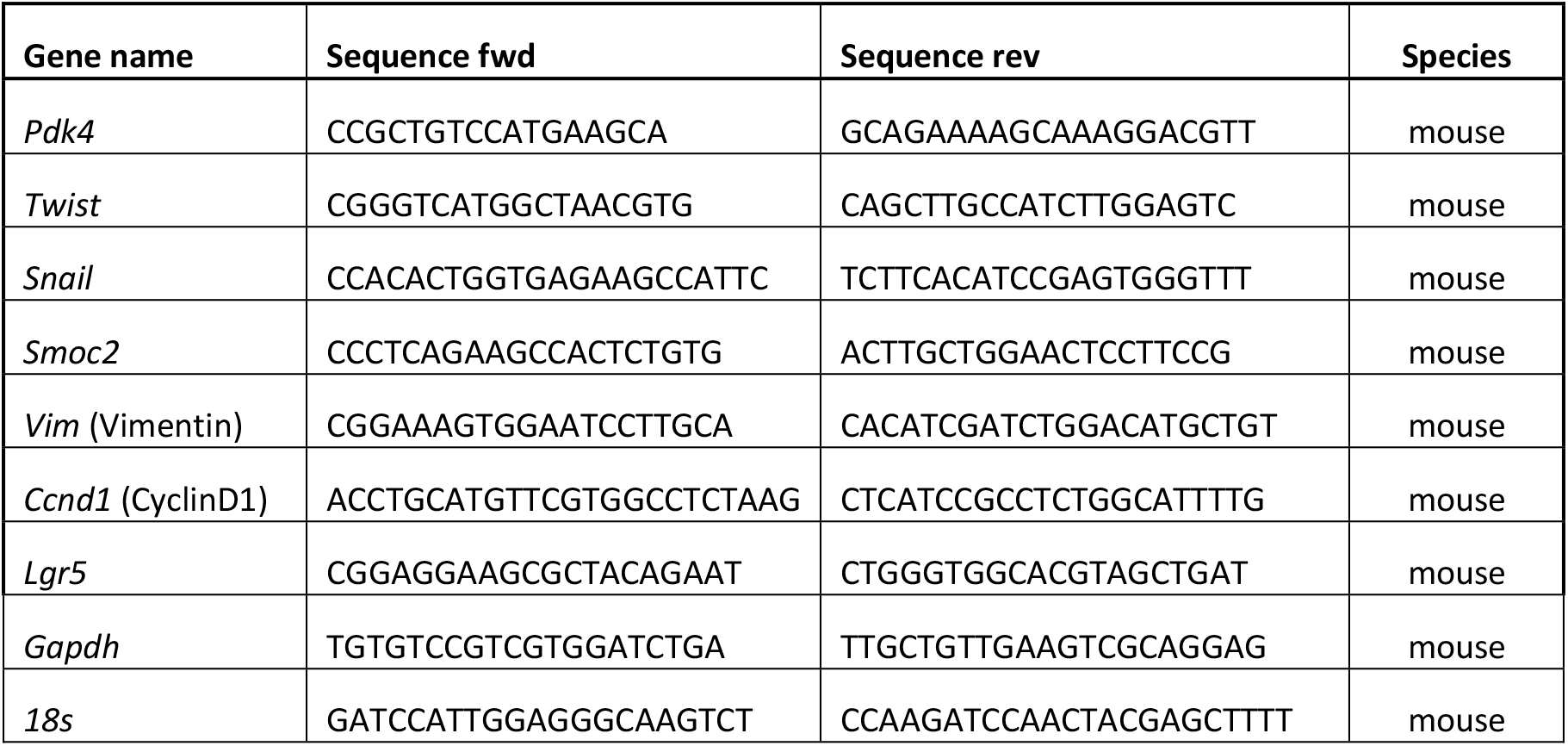
Primers used for qPCR analysis

### Immunofluorescence

Freshly dissected colon tissue was swiss-rolled and placed in 4% PFA overnight. Samples were paraffin embedded and 5 uM sections cut. Paraffin sections were rehydrated and washed with PBS. Then, tissues were incubated with 1% BSA/PBS supplemented with 5% normal serum donkey serum to quench the nonspecific binding of antibodies. Goat anti-E-cadherin (R&D Systems, 1:200) was incubated overnight, washed, secondary stain with anti-goat IgG 633 (LifeTechnologies) and counterstained with DAPI. Samples were imaged using Leica AF6000 microscope.

### GC-MS analysis of metabolites

Extraction and measurement of metabolites were previously described (*50*) with some modifications. Tumor and normal intestinal tissues and tumor organoids (approximately 10 mg) were added to 125 μl methanol, 150 μl Milli-Q water containing internal standard (100 μmol/l 2-isopropylmalic acid) and 60 μl CHCl3 and disrupted with zirconia beads using Micro Smash MS-100 (Tomy Seiko). All samples were shaken at 1,200 rpm for 30 min at 37°C. After centrifugation at 16,000 × g for 5 min at room temperature, 250 μl of the supernatant were transferred to a new tube and 200 μl of Milli-Q water added. After being mixed, the solution was centrifuged at 16,000 × g for 5 min at room temperature, and 250 μl of the supernatant were transferred to a new tube. Samples were evaporated for 20 min at 40°C, and then lyophilized using a freeze dryer. Dried extracts were firstly methoxymated with 40 μl of 20 mg/ml methoxyamine hydrochloride (Sigma-Aldrich) dissolved in pyridine. After adding the derivatization agent, samples were shaken at 1,200 rpm for 90 min at 30°C. Samples were then silylated with 20 μl of N-methyl-N-trimethylsilyl-trifluoroacetamide (GL Science) for 30 min at 37°C with shaking at 1,200 rpm. After derivatization, samples were centrifuged at 16,000 × g for 5 min at room temperature, and the supernatant transferred to glass vial for gas chromatography-tandem mass spectrometry measurement using a GCMS-TQ8030 triple quadrupole mass spectrometer (Shimadzu) with a capillary column (BPX5, SGE Analytical Science). The GC program was previously described (*50*). Data processing was performed using LabSolutions Insight (Shimadzu).

The quantified metabolome data was statistically analyzed using Orthogonal Partial Least Square-Discriminant Analysis (OPLS-DA) with SIMCAP+ software (Version 12.0.1.0, Umetrics, Umeå, Sweden) using UV scaling method. Potential metabolites were selected based on the Variable Importance in Projection (VIP) score greater than 1.0 and uploaded to MetaboAnalyst 3.0 (http://www.metaboanalyst.ca) for pathway analysis. The functional pathway analysis of potential biomarkers was based on the database source of the Kyoto Encyclopedia of Genes and Genomes (http://www.genome.jp/kegg/).

### Tumor organoid establishment and culture

Tumor organoids were established as previously described (*51*), with some alterations. Buffers and culture medium components are listed in Table 2. Tumors were dissected from the colons of mice that had CAC-induced tumors and from the SI and colon of *Apc^min/+^* mice. Tumors were washed in cold PBS then incubated in 10 mL chelation buffer with 2mM EDTA for 60 mins on ice. Tubes were shaken vigorously by hand (removes most normal epithelium) and EDTA buffer was removed and tumor tissue then washed 3x with chelation buffer. Tissue was then incubated in digestion buffer for 30 minutes at 37°C with shaking. Supernatant was filtered through a 70 μM filter, remaining tissue wash once with basic medium and also filtered. Cells were then pelleted (300 g for 3 minutes) washed once then resuspended in Matrigel (Corning 356231) (50 uL per well) and cultured in 24-well plate with 500 μL complete medium. Once established for 1 to 2 weeks, cultures were switched to EGF-only medium.

**Table 2:** Intestinal tumor organoid culture reagents

|  |  |
| --- | --- |
| **Chelation buffer** | in distilled water:<br>- 5.6 mmol/L Na<sub>2</sub>HPO<sub>4</sub><br>- 8.0 mmol/L KH<sub>2</sub>PO<sub>4</sub><br>- 96.2 mmol/L NaCl<br>- 1.6 mmol/L KCl<br>- 43.4 mmol/L sucrose<br>- 54.9 mmol/L D-sorbitol<br>- 0.5 mmol/L DL-dithiothreitol (added fresh) |
| **Digestion buffer** | DMEM:<br>- 2.5% FBS<br>- pen/strep<br>- 400U/mL collagenase D<br>- 25 U/mL Dispase |
| **Basic medium** | - Advanced DMEM F12 – 500mL (Life Technologies)<br>- Pen/Strep 1:100<br>- Glutamax – 1:100<br>- HEPES 1:100 |
| **2xN2/B27/NA medium** | - N2 (200 uL of 100x)<br>- B27 (400 uL of 50x)<br>- NA (50 uL of 500 mM)<br>- Added to 10 mL of Basic medium – use within 1-2 weeks |
| **Complete medium** | - Basic medium + 2xN2/B27/NA (1:1)<br>- mNoggin. Peprotech Cat#250-38. 1000x stock: 100 ug/mL in 0.1% BSA/PBS. 20 uL aliquots. Store at -20.<br>- mEGF. Peprotech Cat#315-09. 10,000x stock: 500 ug/mL 0.1% BSA/PBS. 5uL aliquots. *before use, add 45uL basic medium to make 1000x solution.<br>- Rspol conditioned medium (CM) 1:20<br>- CHIR99021 (GSK3 inhibitor). 5mM in DMSO.<br>- Y-27632 (ROCK inhibitor). 10mM in PBS. Use only for recovering frozen stock, or first establishing.<br>Make as you need it, use immediately |
| **Tumor medium** | - Basic medium + 2xN2/B27/NA (1:1)<br>- mEGF (1:1000)<br>- ROCK inhibitor (Y27632) 1:500 for recovery (not in growth)<br>Make as you need it, use immediately |

For maintenance, organoids were split every week. In brief, all tubes and tips were pre-coated with FBS, organoids were manually disrupted from the Matrigel using P500 pipette and transferred to a 50 mL Falcon tube. Tubes were centrifuged at 300 g for 5 mins at 4 deg. Excess medium was removed carefully to not removed Matrigel containing the cell pellet. Organoids were resuspended in 4 mL Basic medium, transferred to a 15 mL tube and pipetted thoroughly using a fire-polished glass pipette to break up the organoids. These were centrifuged and resuspended in Matrigel and plated out. Split ratios of around 1:4 to 1:6 depending on density.

### *STm*^Δ*aroA*^ infection of tumor organoids

Tumor organoids were infected at day 5 post-split to ensure good organoid size and integrity. *STm*^Δ*aroA*^ was grown overnight and sub-cultured as described above. 5 μL containing 1×10^8^ CFU was dropped into the culture medium (or PBS control) and left for 2 hrs to allow for bacterial invasion of the Matrigel and organoids. After 2 hrs medium was removed and Matrigel washed 2x with PBS and medium replaced and gentamycin added. These were cultured overnight and organoids collected 24 hrs after initial infection for analysis. For qPCR analysis, buffer RLT was added directly to the culture plate (after removing medium and washing with PBS) which completely dissolved the Matrigel, these were then processed for RNA isolation as described above. For metabolome and succinate assays, culture medium was removed and the Matrigel washed with PBS. BD Cell Recovery Solution was added and plate kept on ice for 1 hr to dissolve the Matrigel. Organoids were collected into Eppendorf tubes spun and washed twice with cold PBS. Organoids were then snap frozen and stored at −20 until analysis.

Heat killed *STm*^Δ*aroA*^ was prepared by incubating at 95°C for 5 minutes. Effective killing was tested by plating out on LB agar. SN was prepared by growing *STm*^Δ*aroA*^ in tumor culture medium until an OD_600_ of approximately 0.7. SN was filtered with a 10 kDa cut-off columns. 1×10^5^ heat-killed *STm*^Δ*aroA*^ were used as this is the average CFU count obtained from the live infections after 24 hours, and the amount of SN added was also calculated based on this CFU.

### Succinate Assay

Succinate assay (Sigma) was performed on cell lysates as per the manufacturers protocol and 96-well plates were measured on a spectrophotometer (Spectrostar Nano – BMG Labtech) at the respective wavelengths.

## Acknowledgments

Thank you to David Bending, Haydn Munford, Adam Cunningham, Jeff Cole and Nobuo Sasaki for critical reading of the manuscript and suggestions.

## Author contributions

KMM conceived the project, designed and performed experiments throughout, analyzed data and wrote the manuscript. MT provided technical assistance throughout and maintained animal breeding. YN performed the GC-MS metabolomics experiments and analyzed data. GMM, IE and AC performed organoid experiments. HOd analysed RNAseq data. TK generated mCherry and GFP-expressing *STm*^Δ*aroA*^, helped establish organoid expertise, and sought local experimental ethics authority. HO advised on the project, manuscript, and provided funding.

## Data and materials availability

RNAseq data is uploaded and available online (Gene Expression Omnibus: GSE136029).

## Supplementary Materials

**Figure S1.**
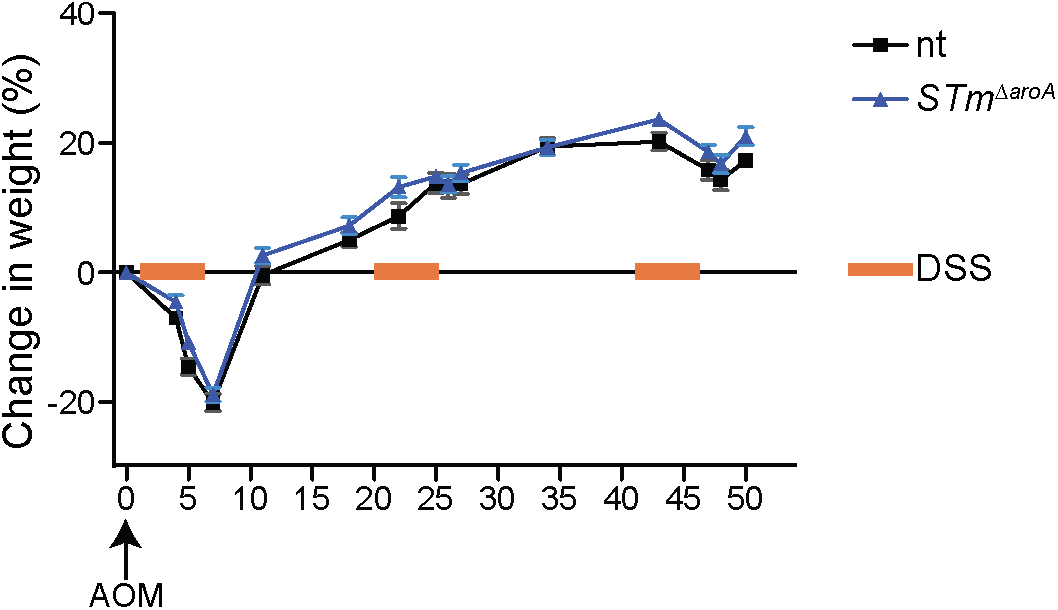
Mouse weight change during AOM/DSS CAC induction. Mice were treated with AOM and DSS (See Figure 1A and methods) to induce colitis-associated cancer. Mouse weight was monitored over the induction period and mice allocated to treatment or non-treatment groups after the last dose of DSS, assuring similar DSS-induced weight change between groups;. Mice represented here are the same group in Figure IB.

**Figure S2:**
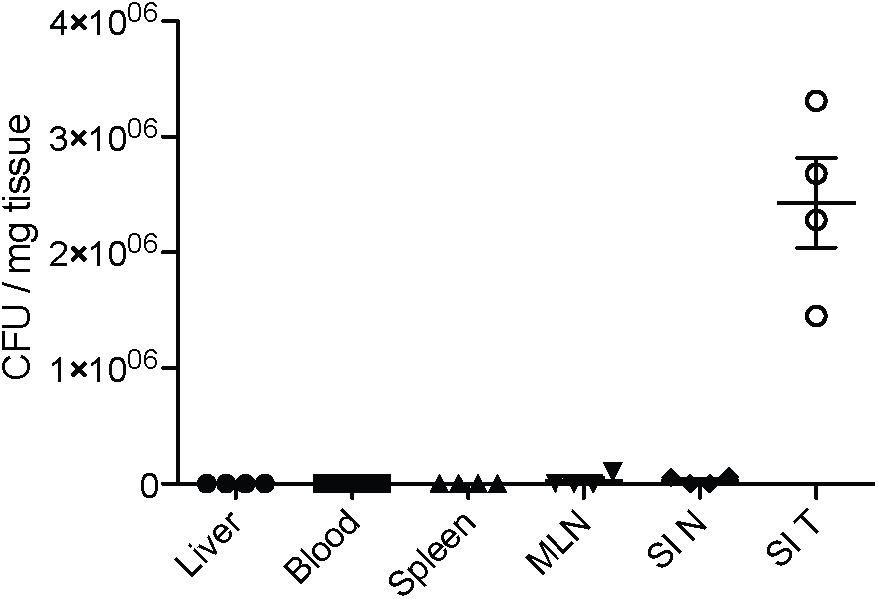
CFU in peripheral organs and SI 24 hours after *STm*^Δ*aroA*^ treatment. *Apc^min/+^* mice were given 5×109 CFU of STmAaroA by oral gavage and bacterial colonization was assessed 24 hours later. Blood, livei, spleen, mesenteric lymph nodes (MLN), normal (N) small intestinal (SI) tissue and SI tumors were dissected, homogenized in PBS −0.1% triton-X and plated onto LB agar plates containing ampicillin.

**Figure S3:**
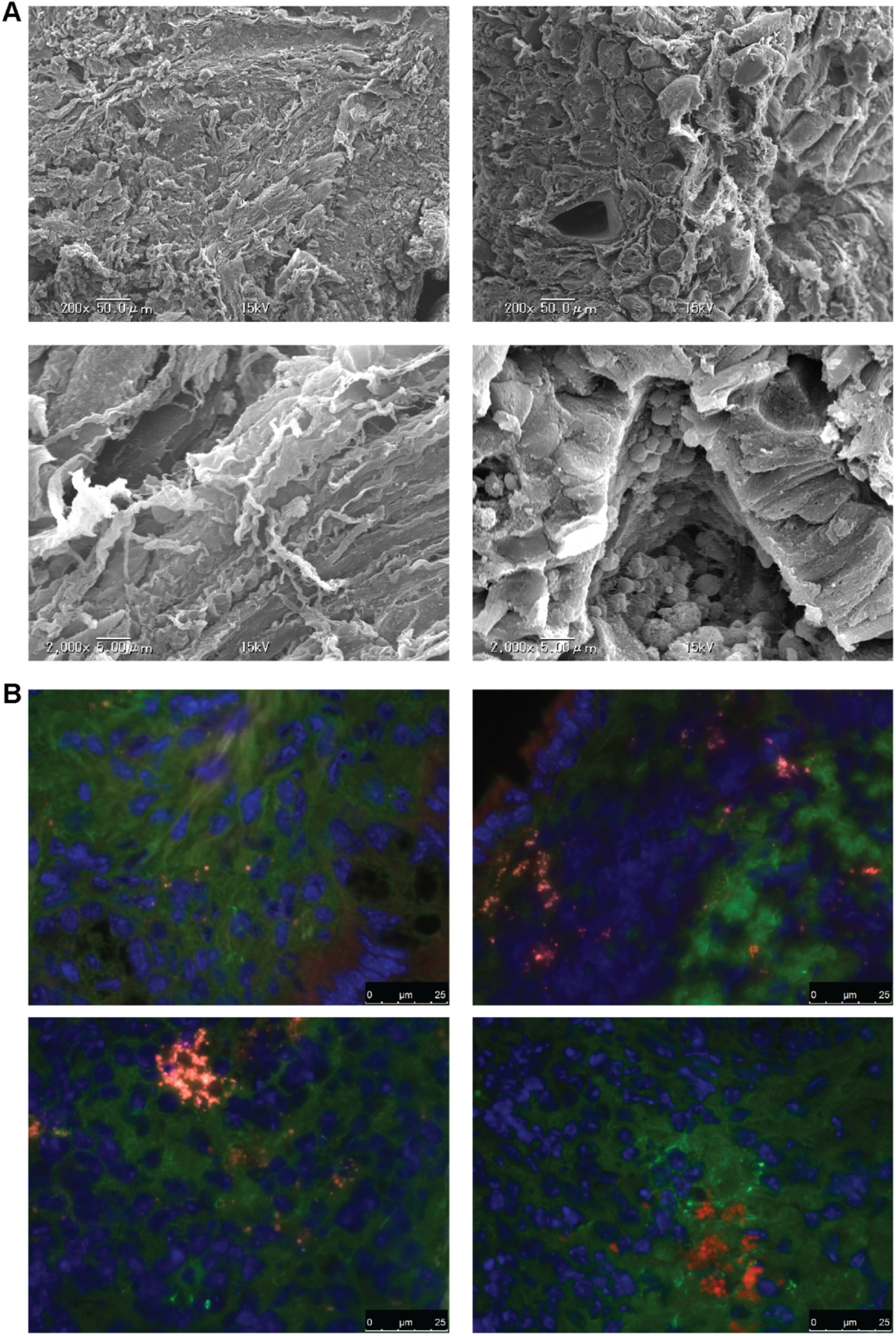
Scanning electron microscopy of and immunohistochemistry of tumors. **(A)** Mice bearing CAC colon tumors were given control vehicle by oral gavage and tissues were taken 24 hours later and prepared for SEM. **(B)** CAC tumor-bearing mice were treated for 6 weeks with mCherry-expressing STm^ΔaroA^. Tissue sections were prepared for IHC. Red = STim^ΔarnA^, Green = F-actin, Blue = DAPI.

**Figure S4:**
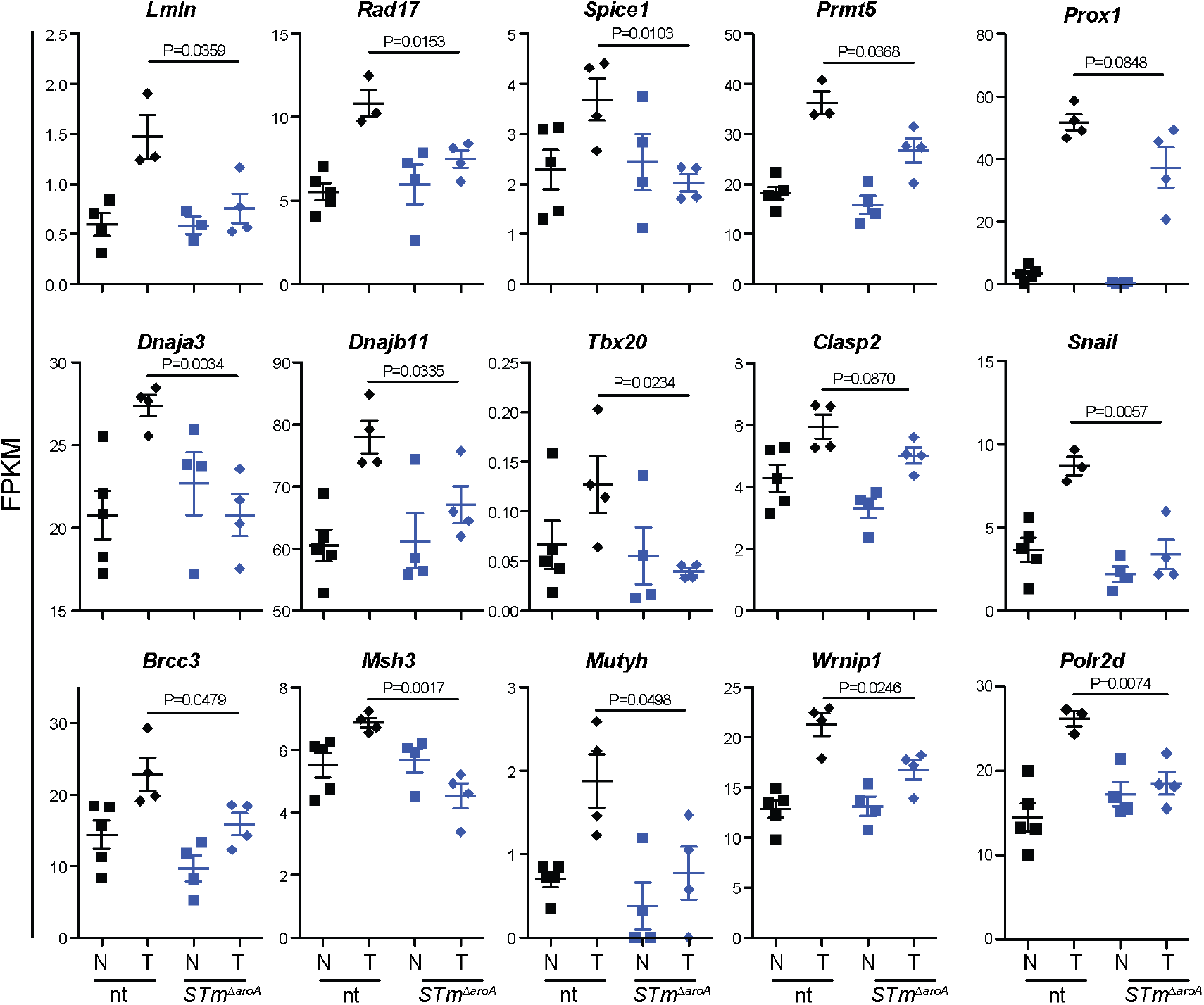
*STm^ΔaroA^* treatment alters the transcriptional landscape of tumors. A selection of differentially expressed genes (DEGs) identified by RNA sequencing. Tumors shows an increased expression of a range of gene associated with processes from cell cycle regulation, DNA repair to epithelial-to-mesenchymal transition. Many of these are reduced following STmAaroA treatment.

**Figure S5:**
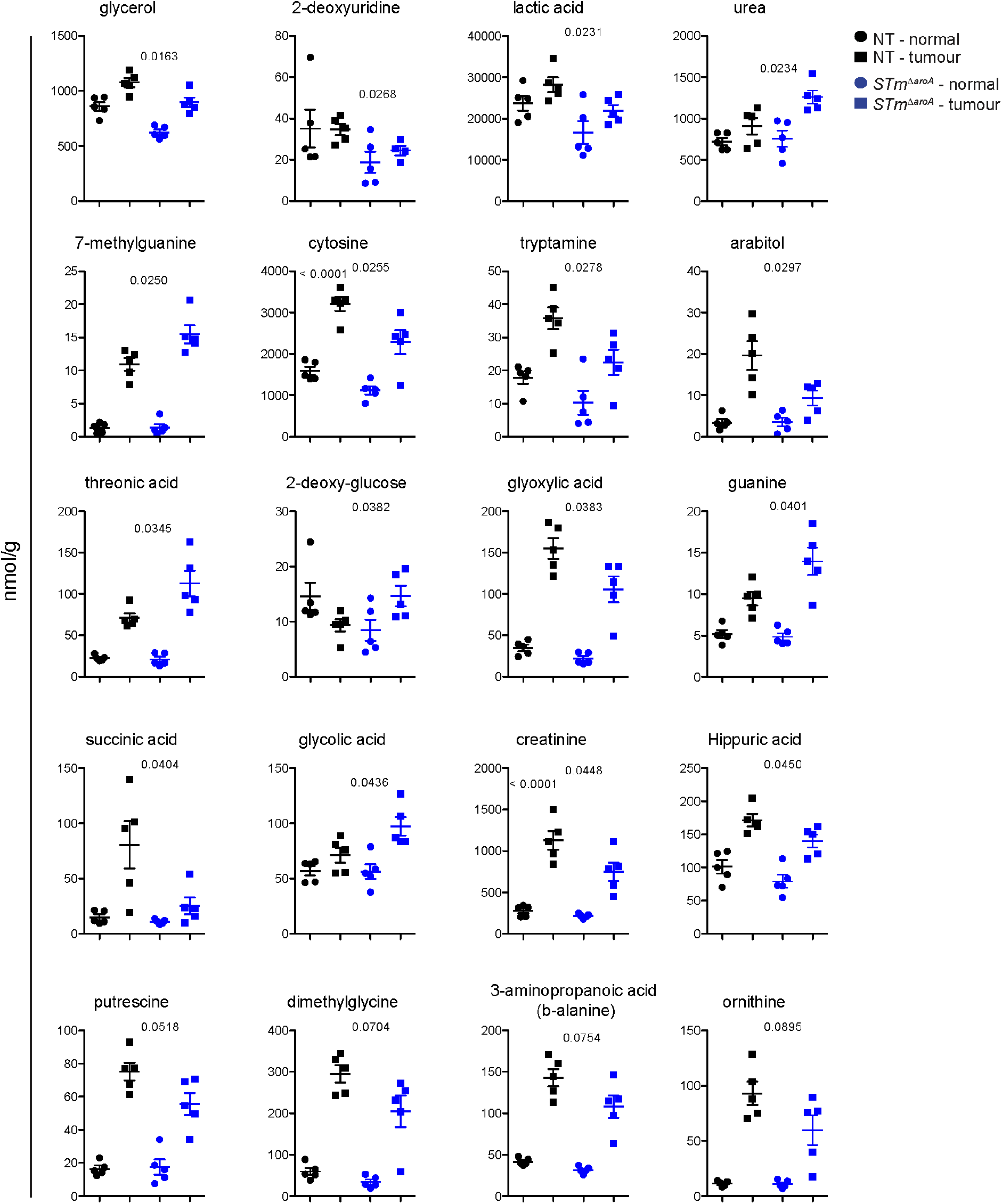
Metabolomics analysis after 6 weeks *STm*^Δ*aroA*^ treatment *in vivo*. Top altered metabolites detected as described in figure 5.

**Figure S6:**
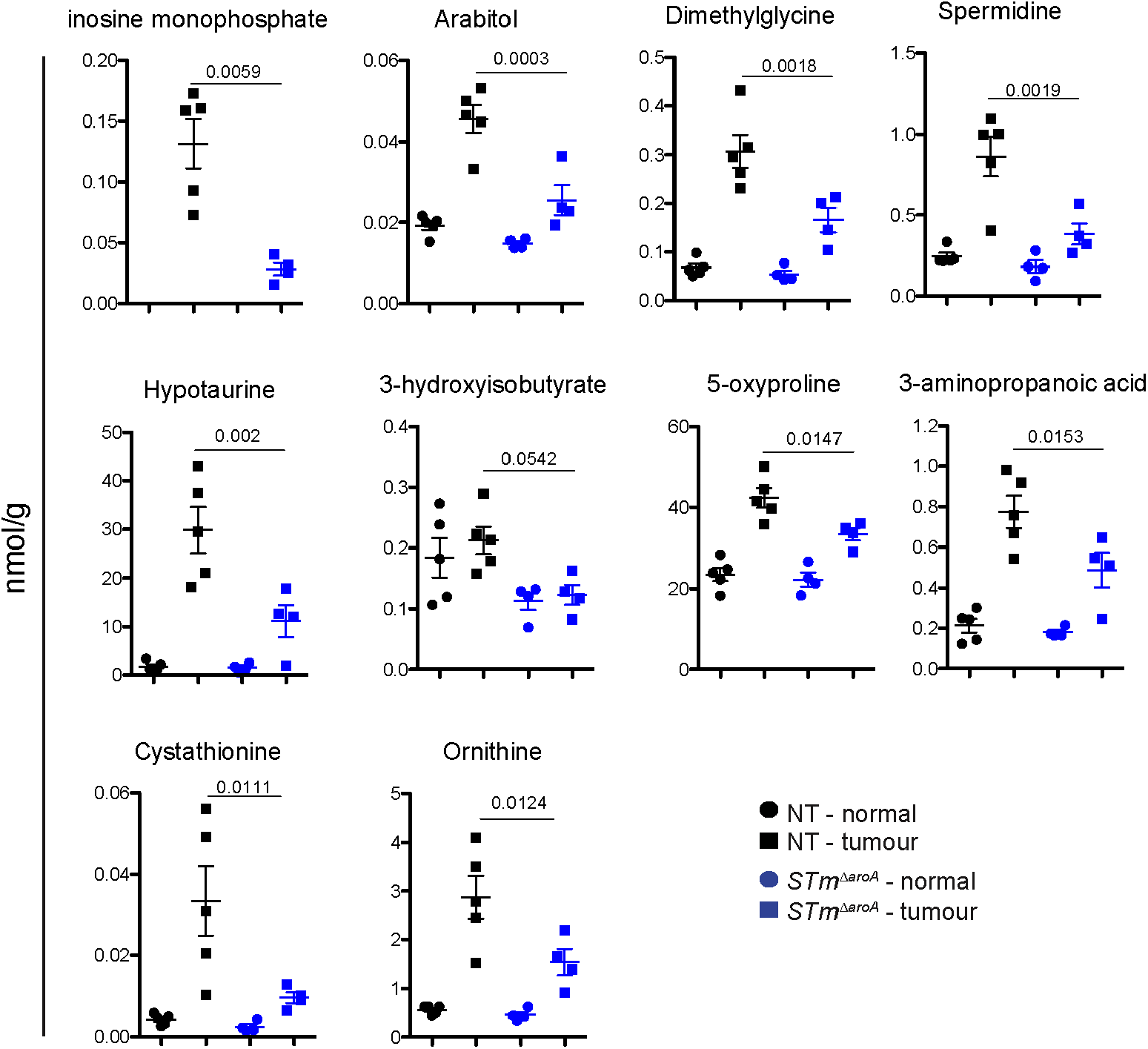
Metabolomics analysis after 24 hours *STm*^Δ*aroA*^ treatment *in vivo*. Top altered metabolites detected as described in figure 5.

**Figure S7:**
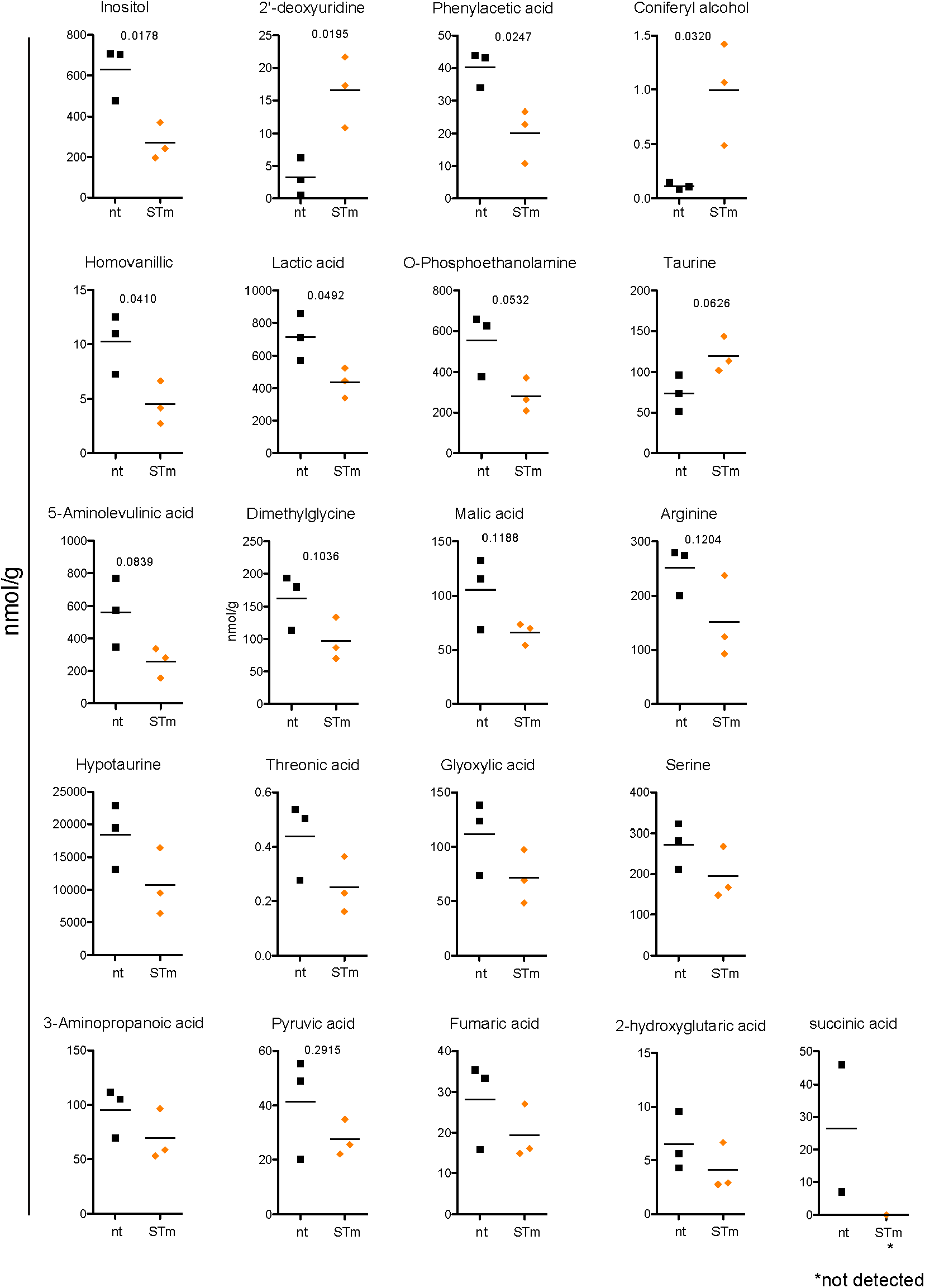
Metabolomics analysis after 24 hours *STm*^Δ*aroA*^ treatment *in vivo*. Top altered metabolites detected as described in figure 5.

**Figure S8:**
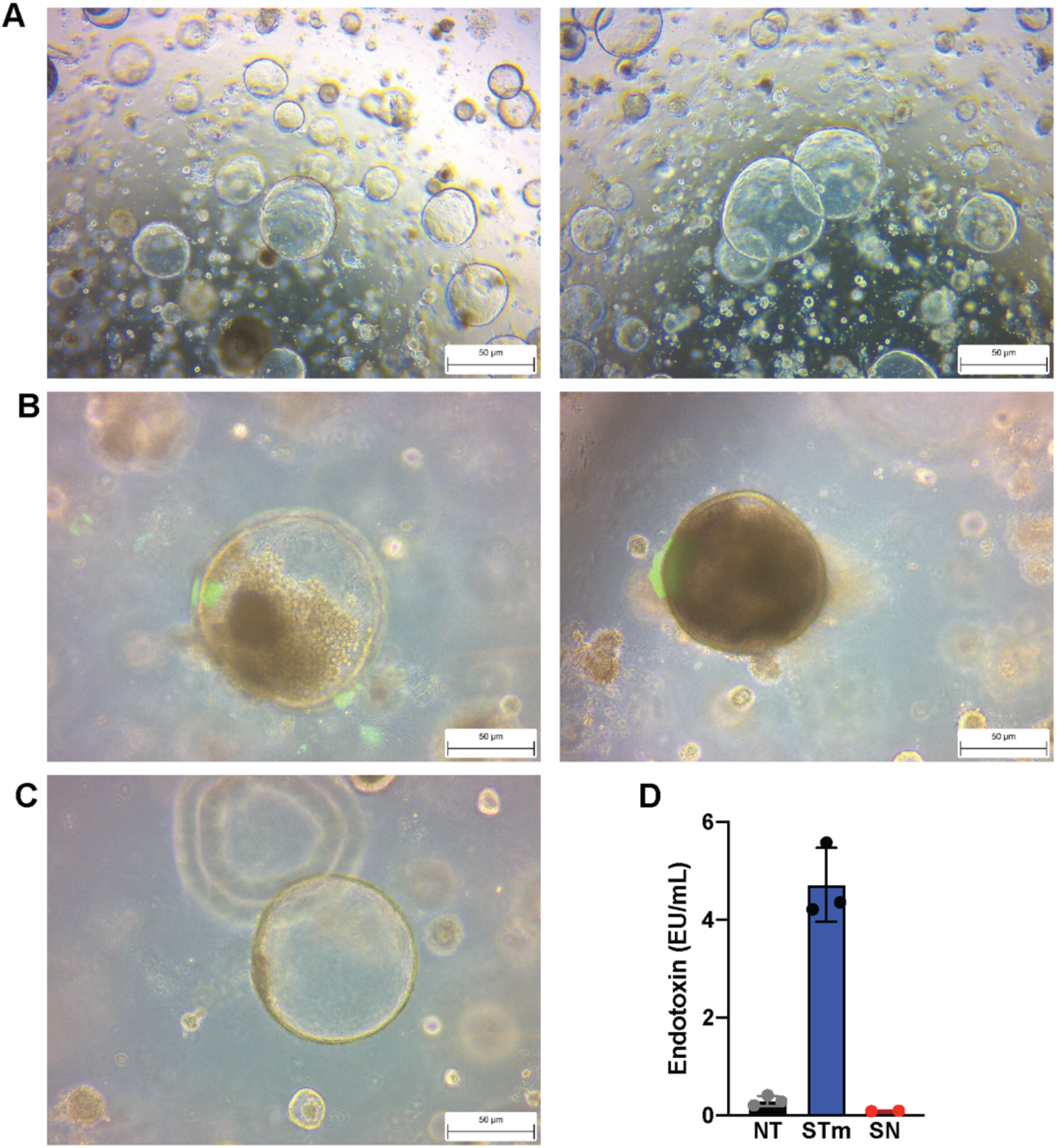
Tumor organoid morphology and infection. **(A)** Tumor organoids are grown as indicated in methods. Representative images of *Apc^min^+*-derived tumor organoids at D5 post-split before infection. **(B)** 24 hours post-infection with GFP-expressing *STm*^Δ*aroA*^, brightfield overlayed with GFP, **(C)** 24 hours control treated, brightfield overlayed with GFP

**Table S1:** List of metabolites with VIP score > 1 in tumors treated with *STm*^Δ*aroA*^ for 6 weeks.

| Name | VIP score | fold change | P value |
| --- | --- | --- | --- |
| Glycerol | 1.9801 | 0.8329 | 0.0163 |
| 2'-Deoxyuridine | 1.9727 | 0.7054 | 0.0268 |
| Lactic acid | 1.9058 | 0.7767 | 0.0231 |
| Urea | 1.9031 | 1.3869 | 0.0234 |
| 7-Methylguanine | 1.8888 | 1.4163 | 0.0250 |
| Cytosine | 1.8837 | 0.7140 | 0.0255 |
| Tryptamine | 1.8638 | 0.6275 | 0.0278 |
| Arabitol | 1.8477 | 0.4773 | 0.0297 |
| Threonic acid | 1.8113 | 1.5825 | 0.0345 |
| 2-Deoxy-glucose | 1.7847 | 1.5725 | 0.0382 |
| Glyoxylic acid | 1.7838 | 0.6815 | 0.0383 |
| Guanine | 1.7717 | 1.4754 | 0.0401 |
| Succinic acid | 1.7699 | 0.3128 | 0.0404 |
| Glycolic acid | 1.7492 | 1.3657 | 0.0436 |
| Creatinine | 1.7423 | 0.6650 | 0.0448 |
| Hippuric acid | 1.7407 | 0.8170 | 0.0450 |
| Putrescine | 1.6135 | 0.7501 | 0.0691 |
| Dimethylglycine | 1.6076 | 0.6924 | 0.0704 |
| 3-Aminoisobutyric acid | 1.5878 | 1.9660 | 0.0748 |
| 3-Aminopropanoic acid | 1.5855 | 0.7546 | 0.0754 |
| Xylose | 1.5652 | 2.2137 | 0.0801 |
| Ornithine | 1.5272 | 0.6445 | 0.0895 |
| 5-Aminovaleric acid | 1.5222 | 4.8578 | 0.0908 |
| Indol-3-acetic acid | 1.4590 | 1.6265 | 0.1081 |
| Cellobiose | 1.4068 | 1.6114 | 0.1238 |
| Xanthosine | 1.3765 | 1.7125 | 0.4535 |
| 3-Hydroxyglutaric acid | 1.3594 | 0.8171 | 0.1393 |
| Malic acid | 1.3564 | 0.8042 | 0.1403 |
| Glycyl-Glycine | 1.3298 | 0.8518 | 0.1495 |
| Sarcosine | 1.3274 | 2.5237 | 0.1504 |
| Allantoin | 1.3171 | 1.3080 | 0.1540 |
| Methionine | 1.2888 | 1.6375 | 0.1644 |
| Galactosamine | 1.2835 | 0.7079 | 0.1664 |
| Galactose | 1.2715 | 0.7943 | 0.1710 |
| Psicose | 1.2256 | 0.7198 | 0.1891 |
| Azelaic acid | 1.2253 | 2.3617 | 0.1892 |
| Norepinephrine | 1.2137 | 1.4135 | 0.1940 |
| Maltose | 1.1786 | 0.6873 | 0.2087 |
| Thymine | 1.1628 | 0.7944 | 0.2156 |
| O-Acetylserine | 1.1543 | 0.7792 | 0.2193 |
| Fucose | 1.1515 | 1.5400 | 0.2206 |
| Glucono-1,5-lactone | 1.1514 | 0.8112 | 0.2206 |
| Mannose | 1.1222 | 0.8146 | 0.2338 |
| Dihydrouracil | 1.1212 | 0.3632 | 0.2527 |
| Lyxose | 1.1023 | 1.2183 | 0.2430 |
| Lactitol | 1.1004 | 0.1406 | 0.2469 |
| 3-Hydroxybutyric acid | 1.0600 | 0.7720 | 0.2634 |
| Fructose | 1.0328 | 0.8219 | 0.2769 |
| Phosphoric acid | 1.0182 | 0.9116 | 0.2843 |
| Hypotaurine | 1.0104 | 0.6991 | 0.2883 |
| 2-Aminoadipic acid | 1.0064 | 1.3342 | 0.2904 |

**Table S2:** List of metabolites with VIP score > 1 in tumors treated with *STm*^Δ*aroA*^ for 24 hours in vivo.

| Metabolite | VIP score | fold change | P value |
| --- | --- | --- | --- |
| Inosine monophosphate | 1.9117 | 0.2165 | 0.0031 |
| Ribitol | 1.9078 | 0.5208 | 0.0032 |
| Cellobiose | 1.8644 | 0.3229 | 0.0049 |
| Arabitol | 1.8548 | 0.5605 | 0.0054 |
| Citric acid | 1.8209 | 0.6098 | 0.0072 |
| Trehalose | 1.8013 | 0.3828 | 0.0084 |
| Maltose | 1.7993 | 0.4144 | 0.0085 |
| Glucose | 1.7758 | 0.4263 | 0.0101 |
| Fumaric acid | 1.7675 | 0.7185 | 0.0108 |
| Isomaltose | 1.7439 | 0.5221 | 0.0337 |
| Dimethylglycine | 1.7136 | 0.5386 | 0.0154 |
| Lactulose | 1.7134 | 0.6199 | 0.0155 |
| Spermidine | 1.7116 | 0.4437 | 0.0156 |
| Hypotaurine | 1.6867 | 0.3729 | 0.0182 |
| 3-Hydroxyisobutyric acid | 1.6852 | 0.5794 | 0.0184 |
| Lactitol | 1.682 | 0.318 | 0.0187 |
| 5-Oxoproline | 1.6626 | 0.7883 | 0.021 |
| Xanthosine monophosphate | 1.5912 | 0.624 | 0.0307 |
| Xylitol | 1.5887 | 0.4935 | 0.0311 |
| Dihydroxyacetone phosphate | 1.5866 | 0.6799 | 0.0314 |
| Galactitol | 1.5846 | 0.697 | 0.0317 |
| Malic acid | 1.582 | 0.7039 | 0.0321 |
| 3-Aminopropanoic acid | 1.508 | 0.6303 | 0.0453 |
| Cystathionine | 1.5071 | 0.2898 | 0.0455 |
| 2-Hydroxyglutaric acid | 1.5053 | 0.653 | 0.0459 |
| Ornithine | 1.4971 | 0.5365 | 0.0475 |
| Cystine | 1.4846 | 0.1547 | 0.041 |
| Orotic acid | 1.4598 | 0.5003 | 0.0555 |
| O-Phosphoethanolamine | 1.4504 | 0.7158 | 0.0576 |
| Mannitol | 1.4501 | 0.6358 | 0.0577 |
| Succinic acid | 1.4359 | 0.6271 | 0.061 |
| Adenine | 1.4174 | 0.6324 | 0.0656 |
| Galactose | 1.4101 | 0.6707 | 0.0674 |
| Glucono-1,5-lactone | 1.409 | 0.6451 | 0.0677 |
| 4-Hydroxyproline | 1.4049 | 0.655 | 0.0688 |
| Glucosamine | 1.4024 | 1.54 | 0.0694 |
| Glycine | 1.3736 | 0.6654 | 0.0771 |
| 2,3-Bisphosphoglyceric acid | 1.3672 | 1.6316 | 0.0789 |
| Threonine | 1.3359 | 0.7095 | 0.088 |
| Glycerol 3-phosphate | 1.3344 | 4.6072 | 0.0884 |
| Cadaverine | 1.3212 | 0.7518 | 0.0925 |
| Glutamic acid | 1.3211 | 0.7397 | 0.0925 |
| Lactic acid | 1.3108 | 0.8793 | 0.0956 |
| Glycerol 2-phosphate | 1.3081 | 1.7895 | 0.0966 |
| Norvaline | 1.3072 | 0.4694 | 0.0969 |
| Cystamine | 1.2993 | 0.5627 | 0.0994 |
| Threonic acid | 1.2797 | 0.764 | 0.1059 |
| Lysine | 1.2702 | 0.6183 | 0.1091 |
| Phosphoenolpyruvic acid | 1.2513 | 1.3313 | 0.2217 |
| Tryptophan | 1.2467 | 0.4 | 0.1173 |
| Tyramine | 1.242 | 0.6351 | 0.119 |
| Arginine | 1.242 | 0.6349 | 0.119 |
| Ethylmalonic acid | 1.2372 | 1.5596 | 0.1207 |
| Erythrose 4-phosphate | 1.2264 | 0.5996 | 0.1247 |
| Ureidosuccinic acid | 1.2199 | 0.7154 | 0.1271 |
| Niacinamide | 1.2189 | 0.8437 | 0.1275 |
| Histidine | 1.2025 | 0.6424 | 0.1338 |
| Allantoin | 1.1996 | 0.5463 | 0.1349 |
| Proline | 1.1976 | 0.736 | 0.1357 |
| Ribonic acid | 1.1963 | 0.736 | 0.1362 |
| 2-Aminoadipic acid | 1.1784 | 0.6401 | 0.1433 |
| Pyruvic acid | 1.1765 | 0.8362 | 0.1441 |
| 6-Phosphogluconic acid | 1.1526 | 0.4985 | 0.1539 |
| Oleamide | 1.1515 | 0.4691 | 0.1544 |
| Hydroquinone | 1.1409 | 0.67 | 0.1588 |
| 2-Aminoisobutyric acid | 1.1385 | 1.9668 | 0.1599 |
| N-Acetylserine | 1.1154 | 0.7525 | 0.17 |
| Norepinephrine | 1.1102 | 1.2179 | 0.1723 |
| Taurine | 1.1045 | 0.8301 | 0.1748 |
| Mannose 6-phosphate | 1.1003 | 0.7004 | 0.1768 |
| 4-Aminobutyric acid | 1.0925 | 0.789 | 0.1803 |
| Glutamic acid 5-methylester | 1.0873 | 1.3579 | 0.1827 |
| Glucose 6-phosphate | 1.0682 | 0.7218 | 0.1917 |
| Quinolinic acid | 1.0502 | 0.0916 | 0.2003 |
| 2-Phosphoglyceric acid | 1.0498 | 1.6942 | 0.1907 |
| Aspartic acid | 1.0399 | 0.8244 | 0.2054 |
| Allose | 1.0192 | 0.7538 | 0.2158 |
| Glucaric acid | 1.0118 | 0.663 | 0.1895 |
| N-Acetylneuraminic acid | 1.005 | 0.827 | 0.2231 |

**Table S3:** List of metabolites with VIP score > 1 in tumor organoids treated with *STm*^Δ*aroA*^ for 24 hours.

| Metabolite | VIP score | fold change | P value |
| --- | --- | --- | --- |
| Inositol | 1.8382 | 0.4290 | 0.0178 |
| 2'-Deoxyuridine | 1.8269 | 5.1418 | 0.0195 |
| Phenylacetic acid | 1.7964 | 0.4992 | 0.0247 |
| Coniferyl alcohol | 1.7576 | 8.5705 | 0.0320 |
| Homovanillic acid | 1.7155 | 0.4414 | 0.0410 |
| Lactic acid | 1.7002 | 0.6143 | 0.0492 |
| O-Phosphoethanolamine | 1.6650 | 0.5077 | 0.0532 |
| Fucose | 1.6605 | 3.1011 | 0.0543 |
| Indol-3-acetic acid | 1.6476 | 0.7028 | 0.0577 |
| Taurine | 1.6293 | 1.6209 | 0.0626 |
| 5-Aminolevulinic acid | 1.5572 | 0.4599 | 0.0839 |
| Uridine | 1.5306 | 2.1723 | 0.1633 |
| Lyxose | 1.5002 | 0.5528 | 0.1026 |
| Dimethylglycine | 1.4973 | 0.5974 | 0.1036 |
| Galactose | 1.4862 | 2.2448 | 0.1075 |
| Malic acid | 1.4547 | 0.6259 | 0.1188 |
| Arginine | 1.4504 | 0.6035 | 0.1204 |
| Ribonic acid | 1.4259 | 0.2950 | 0.1295 |
| 4-Aminobenzoic acid | 1.4205 | 0.5189 | 0.1316 |
| Hypotaurine | 1.4123 | 0.5822 | 0.1348 |
| Threonic acid | 1.4033 | 0.5742 | 0.1383 |
| Mannitol | 1.3841 | 3.2098 | 0.1437 |
| Spermidine | 1.3798 | 3.1311 | 0.1476 |
| Glyoxylic acid | 1.3191 | 0.6411 | 0.1729 |
| Sebacic acid | 1.3114 | 0.6055 | 0.1762 |
| Guanine | 1.2997 | 2.1673 | 0.1813 |
| Homocysteine | 1.2973 | 2.7287 | 0.1824 |
| 2-Aminoethanol | 1.2938 | 0.7175 | 0.1839 |
| Phosphoenolpyruvic acid | 1.2811 | 1.5814 | 0.1895 |
| Suberic acid | 1.2801 | 0.3291 | 0.1900 |
| Serine | 1.2790 | 0.7142 | 0.1905 |
| Pantothenic acid | 1.2545 | 0.7354 | 0.2016 |
| 3-Aminopropanoic acid | 1.1723 | 0.7260 | 0.2407 |
| Xylose | 1.1591 | 3.2897 | 0.2472 |
| Alanine | 1.1475 | 1.4680 | 0.2530 |
| Arabitol | 1.1097 | 3.7650 | 0.2723 |
| Glucose | 1.1063 | 3.0801 | 0.5069 |
| Sorbitol | 1.1018 | 3.1654 | 0.5090 |
| Tyrosine | 1.0957 | 2.4572 | 0.2795 |
| Pyruvic acid | 1.0729 | 0.6639 | 0.2915 |
| Fumaric acid | 1.0709 | 0.6866 | 0.2926 |
| Dodecanedioic acid | 1.0598 | 0.6416 | 0.2985 |
| 3-Hydroxypropionic acid | 1.0470 | 0.6421 | 0.3054 |
| 2-Hydroxyglutaric acid | 1.0444 | 0.6318 | 0.3068 |
| Ribose | 1.0413 | 1.6181 | 0.3085 |
| Pyridoxamine | 1.0286 | 2.3168 | 0.3154 |
| Glutaric acid | 1.0275 | 0.3534 | 0.3160 |
| Ureidopropionic acid | 1.0198 | 0.5835 | 0.3202 |
| Dopa | 1.0162 | 1.7613 | 0.3222 |
| Adenine | 1.0124 | 1.6859 | 0.3243 |
| Hypoxanthine | 1.0080 | 5.7728 | 0.3267 |
| 4-Aminobutyric acid | 1.0006 | 1.7863 | 0.3308 |

